# Paired aptamer capture and FISH detection of individual virions enables cell-free determination of infectious titer

**DOI:** 10.1101/2022.11.13.516306

**Authors:** Yifang Liu, Jacob L. Potts, Dylan Bloch, Keqing Nian, Caroline A. McCormick, Oleksandra Fanari, Sara H. Rouhanifard

## Abstract

Early detection of viruses can prevent the uncontrolled spread of viral infections. Determination of viral infectivity is also critical for determining the dosage of gene therapies, including vector-based vaccines, CAR T-cell therapies, and CRISPR therapeutics. In both cases, for viral pathogens and viral vector delivery vehicles, fast and accurate measurement of infectious titer is desirable. The most common methods for virus detection are antigen-based (rapid but not sensitive) and reverse transcription polymerase chain reaction (RT-PCR)-based (sensitive but not rapid). Current viral titer methods heavily rely on cultured cells, which introduces variability within labs and between labs. Thus, it is highly desirable to directly determine the infectious titer without using cells. Here, we report the development of a direct, fast, and sensitive assay for virus detection (dubbed rapid-aptamer FISH or raptamer FISH) and cell-free determination of infectious titers. Importantly, we demonstrate that the virions captured are “infectious,” thus serving as a more consistent proxy of infectious titer. This assay is unique because it first captures viruses bearing an intact coat protein using an aptamer, then detects genomes directly in individual virions using fluorescence in situ hybridization (FISH)– thus, it is selective for infectious particles (i.e., positive for coat protein and positive for genome).

## Introduction

Rapid and accurate virus concentration measurement is desirable in both diagnostic and R&D settings. Clinically, rapid detection of infectious viral particles enables a timely diagnosis, and in industry, viral vectors are a significant component of gene therapies, vector-based vaccines, and CAR-T immunotherapies. Currently, the most common methods for quantifying viruses are either based on the enzyme-linked immunosorbent assay (ELISA)-based detection of viral surface antigen^1,2^ which is rapid but not sensitive, or PCR-based detection of the viral genome which is sensitive but not rapid^3,4^. A shortcoming of each of these assays is that presence of either surface antigen, or viral genome does not necessarily indicate the concentration of infectious viral particles.

The infectious titer of a virus is often calculated in transducing units (TU) per mL. The total particle-to-TU ratio is the total number of particles divided by the TUs. For many viruses, the particle-to-TU ratio can be extremely high and variable^5^. For example, the ratio for Varicella-zoster virus is 40,000:1^6^, while for adenovirus (HAdV-C5), it is 58:1^7^. For HIV-1, the ratios have even more variability ranging from 1-10^7^:1^8^. In these cases, the properties measured biochemically may not be directly attributed to those of the infectious particles, thus complicating the quality control and assessment of performance for gene therapies *in vivo* and evaluation of how infectious an individual is at a given moment.

The surface antigen test (ELISA) may detect viruses with surface antigens but no genome. By contrast, PCR-based tests may detect viruses that contain a genome but no surface antigens. For example, the viral RNA of SARS-CoV-2 patients can still be detected even in the absence of an infectious virus^9^. Functional titering of infectious virions is accomplished with a viral plaque assay^10,11^ or tissue culture infectious dose–50 (TCID50)^12^, both cell-based. These methods titer virions by measuring the cytopathology in cell monolayers produced by viral replication. However, functional titering using cell culture can lead to significant variability^13^ from cell type, passage number, and condition of the cells. Therefore, a cell-free titering method is needed and can improve the accuracy and consistency of viral quantification.

Single-molecule RNA FISH (smFISH) is a sensitive method that directly detects RNA. It uses approximately 30 fluorescently labeled oligonucleotide probes to target RNA sequences^14^. The tiling of probes across an RNA sequence amplifies the fluorescent signal locally and enables the direct detection of individual RNA molecules using fluorescence microscopy. SmFISH is frequently used to detect virus-infected cells^15–17^ and tissues^18^. It has also been reported to detect individual virions via the capture of surface antigens ^19–21^; however, these methods required specialized instrumentation (i.e., Raman spectroscopy, TIRF microscopy). Two alternative smFISH methods, TurboFISH^16^ and rvFISH^19–21^ can decrease the total time of the assay from 12 hours to <20 minutes while maintaining the sensitivity and specificity of the assay, making these methods ideal candidates for rapid virus detection. However, TurboFISH detects viral genomes in infected cells, and rvFISH detects all viral particles containing the genome, irrespective of infectivity.

Here we report a rapid assay to directly detect intact virions (dubbed rapid-aptamer FISH or “raptamer FISH”) using epifluorescence microscopy for targeting both viral genomes and coat protein, thus providing a better proxy for the overall infectivity of the detected virus. We apply both antibodies and aptamers that are specific for viral coat proteins to capture virions directly on glass. Once captured, viral genomes were detected by TurboFISH to inform the infectivity of virions (positive for coat protein and positive for genome). The use of two probe sets targeting different genomic regions increases the specificity of viral RNA detection. We demonstrate that infectious virions are bound to the antibodies and aptamers by performing a functional titer on the unbound fraction of the virus and showing a marked decrease in infectivity. Overall, these results demonstrate that raptamer FISH can replace cell-based assays for determining infectivity, providing a useful and fast readout for manufacturers of gene therapies that are delivered via viral vectors, as well as a useful and fast measurement of the infectivity of an individual.

## Results

### Capture and sandwich ELISA detection of recombinant spike protein with anti-spike aptamer

We selected previously reported DNA aptamer sequences (1C and 4C) that were computationally predicted to bind to the receptor binding domain of the SARS-CoV-2 Spike protein^22^. To evaluate the binding properties of the aptamers for our assay, we immobilized aptamers biotinylated on either the 5’ and 3’ ends to streptavidin-coated polystyrene plates (**Figure 1a**). As a positive control, we immobilized biotinylated ACE2 protein to the streptavidin-coated polystyrene plates and used random aptamer sequences as our negative control (**Figure 1b**). We applied recombinant SARS-CoV-2 Spike protein to the immobilized capture reagents and performed a sandwich ELISA using an HRP-conjugated anti-Spike antibody followed by colorimetric detection ^22,23,24^.

**Figure 1:**
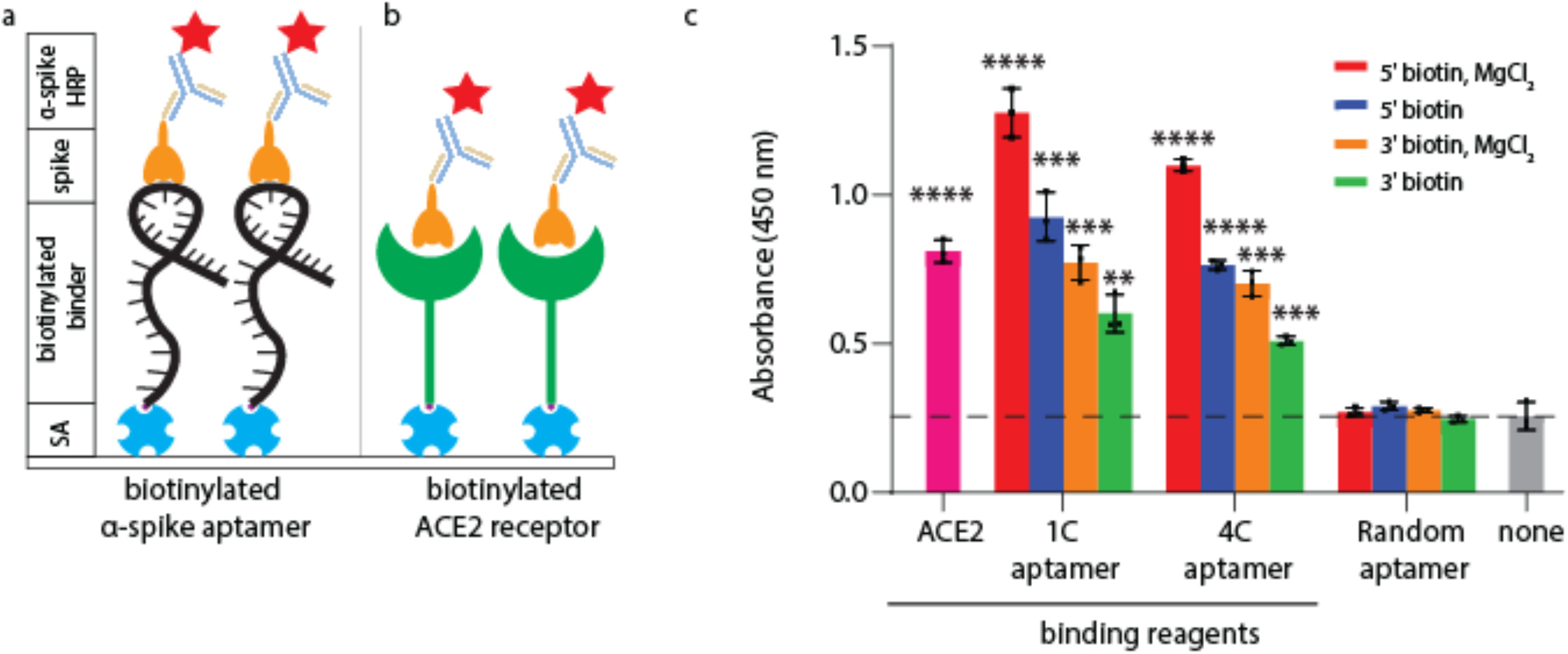
ELISA for capture efficiencies of anti-spike aptamers compared to ACE2 receptor with recombinant SARS-CoV-2 spike protein. a. Aptamers modified with 5’ or 3’ biotin or b. ACE2 proteins with biotin were conjugated to a streptavidin-coated polystyrene plate. Following spike binding (orange), the spike protein was detected with an anti-spike antibody conjugated to HRP, followed by colorimetric detection. c. Measuring the binding specificity of 1C and 4C aptamers, different conditions of aptamers were tested (5’ modified, 3’ modified, ± MgCl_2_. n = 3 biological replicates (mean ± SD). Plates without a biotinylated binder (none) were set as a baseline to show a significant difference from other groups. An unpaired t-test was performed to compare samples. *p < 0.05, **p < 0.01, ***p < 0.001, ****p < 0.0001.

We observed that the 1C aptamers had a stronger affinity towards the spike protein than the 4C aptamers. 5’ biotin-modified 1C had a 4.724±0.079 fold increase of binding and 5’ biotin-modified 4C aptamer had a 4.077±0.083 fold increase of binding as random aptamer in the presence of MgCl_2_. 5’ biotin-modified aptamers captured 1.615±0.055_fold more spike protein than the 3’ modified aptamers. The presence of MgCl_2_ also had a positive effect on the quantity of spike protein captured with a 1.373±0.029 fold increase in absorbance over the condition where MgCl_2_ was omitted, consistent with the previous work ^22,23,24^ (**Figure 1c**). Overall, 5’ biotin-modified 1C aptamers exhibit the highest affinity towards recombinant spike protein with MgCl_2_. This condition was used for the following experiments.

### Capture and detection of spike-pseudotyped lentiviruses with anti-spike aptamer using RT-qPCR

To assess the binding efficiency of virions to the immobilized aptamers, we produced SARS-CoV-2 spike-pseudotyped lentiviral particles according to published work^25^. We immobilized ACE2 and anti-spike aptamer 1C on streptavidin-coated polystyrene plates and exposed them to lentivirus virions to assess the capture efficiency. A random aptamer-coated plate and lentiviral particles pseudotyped with Vesicular Stomatitis Virus Glycoprotein (VSVG)^26^ were used as negative controls to assess the nonspecific binding of the virus to ACE2 and 1C aptamer. The spike-pseudotyped lentivirus and the VSVG-pseudotyped lentivirus were added to the plates at concentrations ranging from 10^5^ to 10^8^ genome copies (determined by RT-qPCR; **Supplementary Figure 1**). After washing, we extracted the RNA from captured virions and performed RT-qPCR targeting the CMV promoter (common between both lentivirus vectors) to determine capture efficiency (**Figure 2a**).

**Figure 2:**
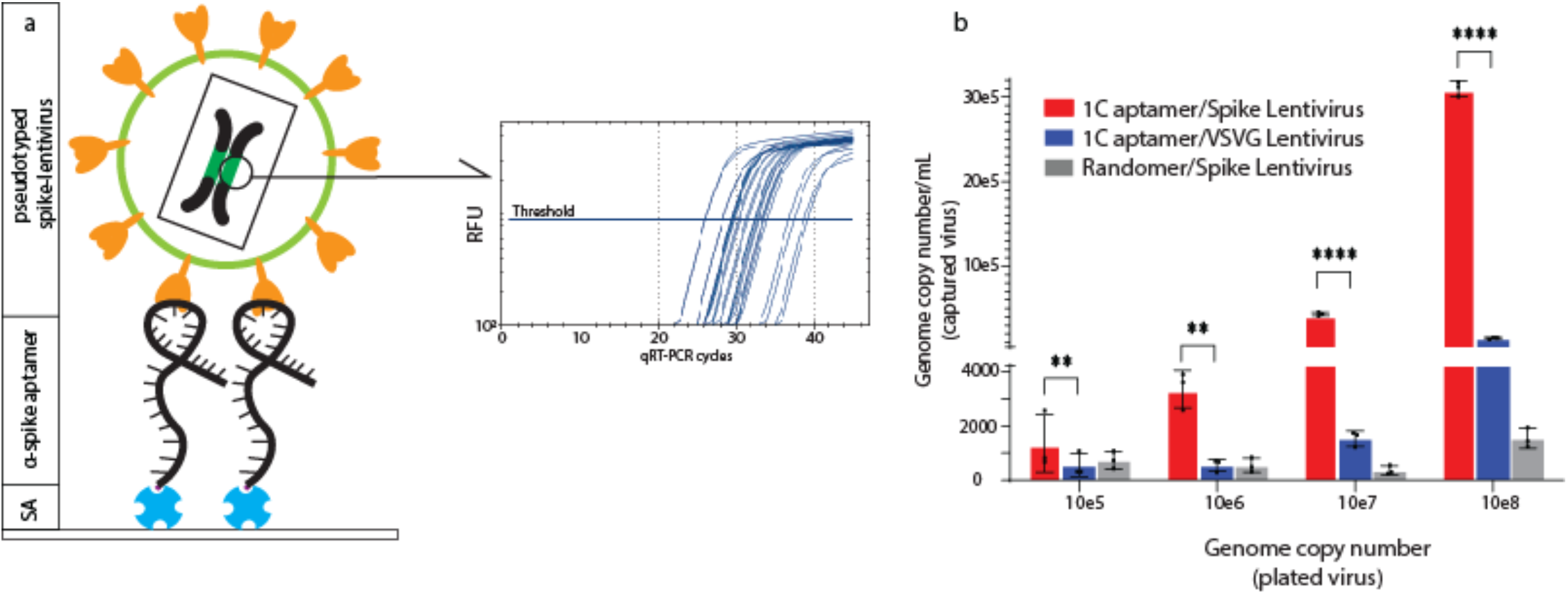
RT-qPCR for capture efficiencies of SARS-COV-2 spike pseudotyped lentivirus by 1C aptamers. a.1C aptamers modified with 5’ biotin were conjugated to a streptavidin-coated polystyrene plate. Following lentivirus capture by the aptamers, RNA was extracted, and RT-qPCR was performed to measure the amount of captured virions. b. Bar graph of genome copies number per mL of spike and VSVG virions captured by 1C aptamers and random aptamers. n = 3 biological replicates (mean ± SD). *p < 0.05, **p < 0.01, ***p < 0.001, ****p < 0.0001.

Specific capture of spike-pseudotyped lentivirus was detected with the 1C capture reagent at a minimum concentration of 10^3^ copies per μl, showing a 2.412-fold increase in genome-containing units over the nonspecific virus captured by the random aptamer. When the concentration was increased to 10^6^ genome-containing units per μl, the capture efficiency increased, showing a 208.292-fold increase in genome copies over the nonspecific virus captured by the random aptamer (**Figure 2b**). The same capture efficiency was also observed in the ACE2 protein (**Supplementary Figure 2**).

We then used a range of aptamer concentrations to coat the streptavidin plate to determine the optimal concentration for virion capture. The capture, RNA extraction, reverse transcription, and qPCR were performed, and six aptamer concentrations ranging from 1 μM- 200 μM were selected. We observed a 45.688-fold higher binding for the 5 μM aptamer concentration over the VSV-G virus and a 10.064-fold higher binding to 200 μM aptamer concentration. We observed that aptamer binding became saturated at around 10 μM (this concentration was used for all following experiments) and a 40-fold increase in spike pseudotyped virus captured over VSV-G virus at 10 μM (**Supplementary Figure 3**).

### Raptamer FISH detection of spike-pseudotyped lentiviruses with 1C aptamer

After validating that our system can capture viruses on polystyrene plates and chambered coverglass (**Supplementary Fig. 4**), we tested the utility of fluorescence *in situ* hybridization (FISH) to detect individual virions that are captured by aptamers in a method dubbed rapid-aptamer FISH (raptamer FISH)(**Figure 3a**). Previous work of detecting virions relies on single molecule FISH (smFISH), which employs hybridization times of up to 16 h^19,27^; however, we applied an alternative method called TurboFISH^16^ that can decrease the hybridization time to 5 minutes to achieve fast detection. We confirmed that the virions would retain their integrity after fixation and permeabilization by capturing the virus and then performing RT-PCR on the captured virions post-methanol fixation and post-TurboFISH (**Supplementary Figure 5**). The chambered coverglass was biotinylated by silane-biotin, and virions were captured as previously described. Briefly, the virus was fixed with methanol after specific capture, and smFISH probes^14^ targeting the SARS-CoV-2 nucleocapsid gene (N gene) were introduced and hybridized. After hybridization, there were further wash steps followed by epifluorescent microscopy (**Figure 3a, 3b, 3e**).

**Figure 3:**
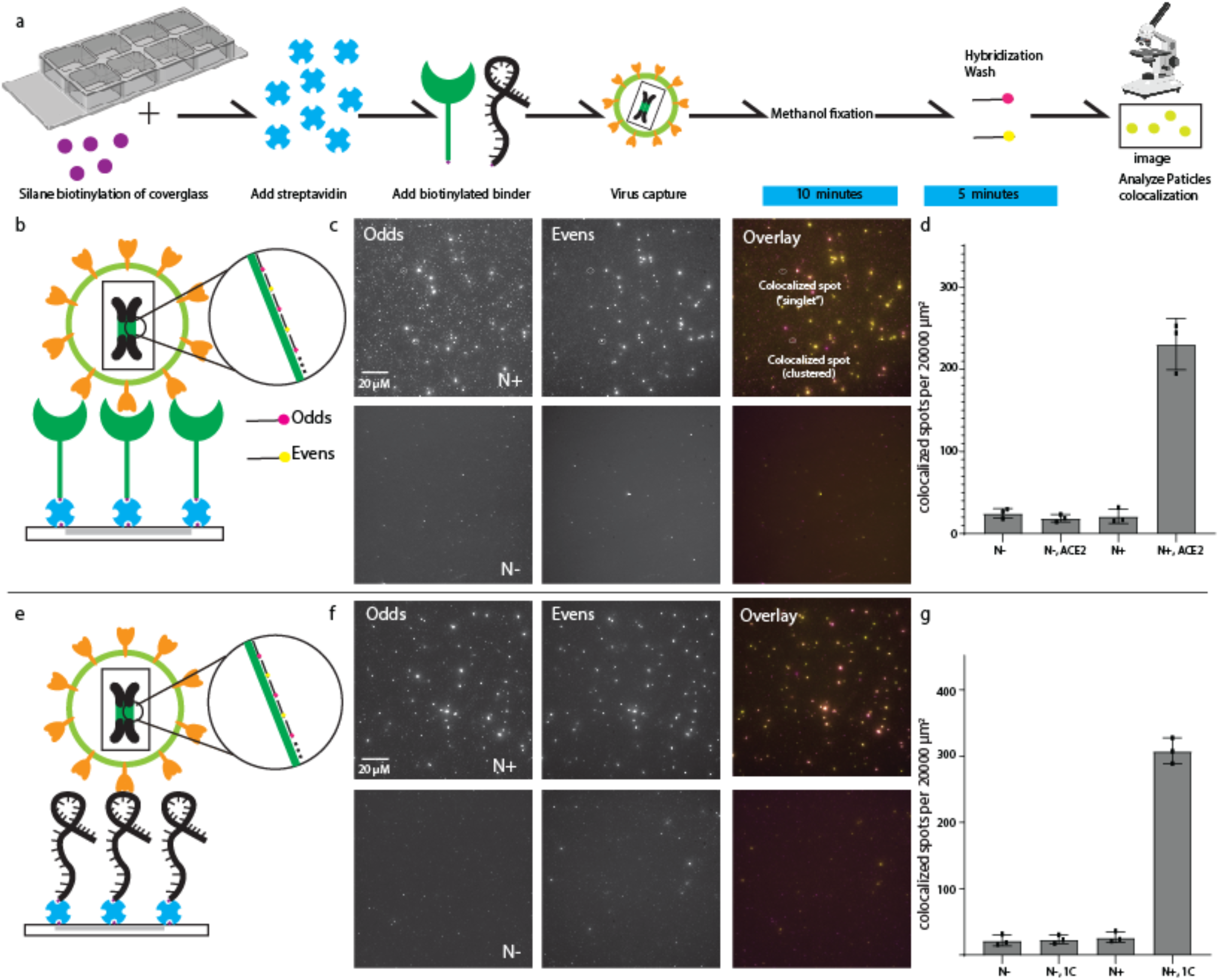
Raptamer FISH detection of individual virions. a. Workflow of raptamer FISH detection. Raptamer FISH using b. ACE2 receptor or e. 1C aptamers. Fluorescent micrographs showing raptamer FISH targeting N gene of the viral genome c, f.. d, g. Colocalized FISH spots per 20000 μm^2^ were quantified in bar graphs by the Matlab colocalization analysis pipeline. 3 biological replicates, 5 images from each replicate (mean ± SD).

The size of the raptamer FISH spots was determined to be 1-500 pixels by particle analysis in Matlab. Lentivirus is known to aggregate^28^. Approximately 79.572% of detected spots were 1-70 pixels in diameter, showing up only in the samples where virus was added. Therefore we reasoned that colocalized spots in the 1-70 pixel range are likely individual virions. We observed that 20.428% of spots were beyond 70 pixels and these large spots were probable viral aggregates (**Supplementary Figure 6a**). The large particles (spots more than 70 pixels) showed the same colocalization rates as the small particles, indicating that the viral aggregates were infectious (**Supplementary Figure 6b**).

To confirm the detection of a true viral particle, we used two sets of probes (odds and evens) to mitigate the detection of false positive spots^14^. The specificity of probes for N gene was demonstrated previously^18^. To confirm specific viral detection, we applied raptamer FISH to a series of negative controls, including the slides not treated with ACE2 protein and the slides not treated with 1C aptamers. Additionally, we tested viruses whose genome contains the N gene (N+) or not (N-) (**Figure 3b-g**).

For the functionalized slides treated with ACE2 or 1C aptamer and in the presence of N+ virus, we observed a 10.755-fold and 12.780-fold increase of colocalized spots, respectively, as compared to the samples without binding agents and N gene (**Figure 3b-g**).

### Simultaneous raptamer FISH detection of multiple virus strains

To test whether the raptamer FISH system can detect multiple strains, we performed the assay on a mixed population of spike coated virions containing two distinct genomes. N+ virions, as described previously, were mixed in different ratios with virions containing the luciferase gene (N-). The N gene probe targets the N gene of spike-coated lentivirus (S+N+), and the luciferase probe targets the luciferase of spike-coated lentivirus without the N gene (S+N-) was combined to detect two types of virions simultaneously. Two sets of probes (odds and evens) with two dyes for each target were used to detect the colocalized FISH spots (**Figure 4a**). Virions were mixed with different concentration ratios, then captured, fixed, and detected by TurboFISH simultaneously (**Figure 4b**).

**Figure 4:**
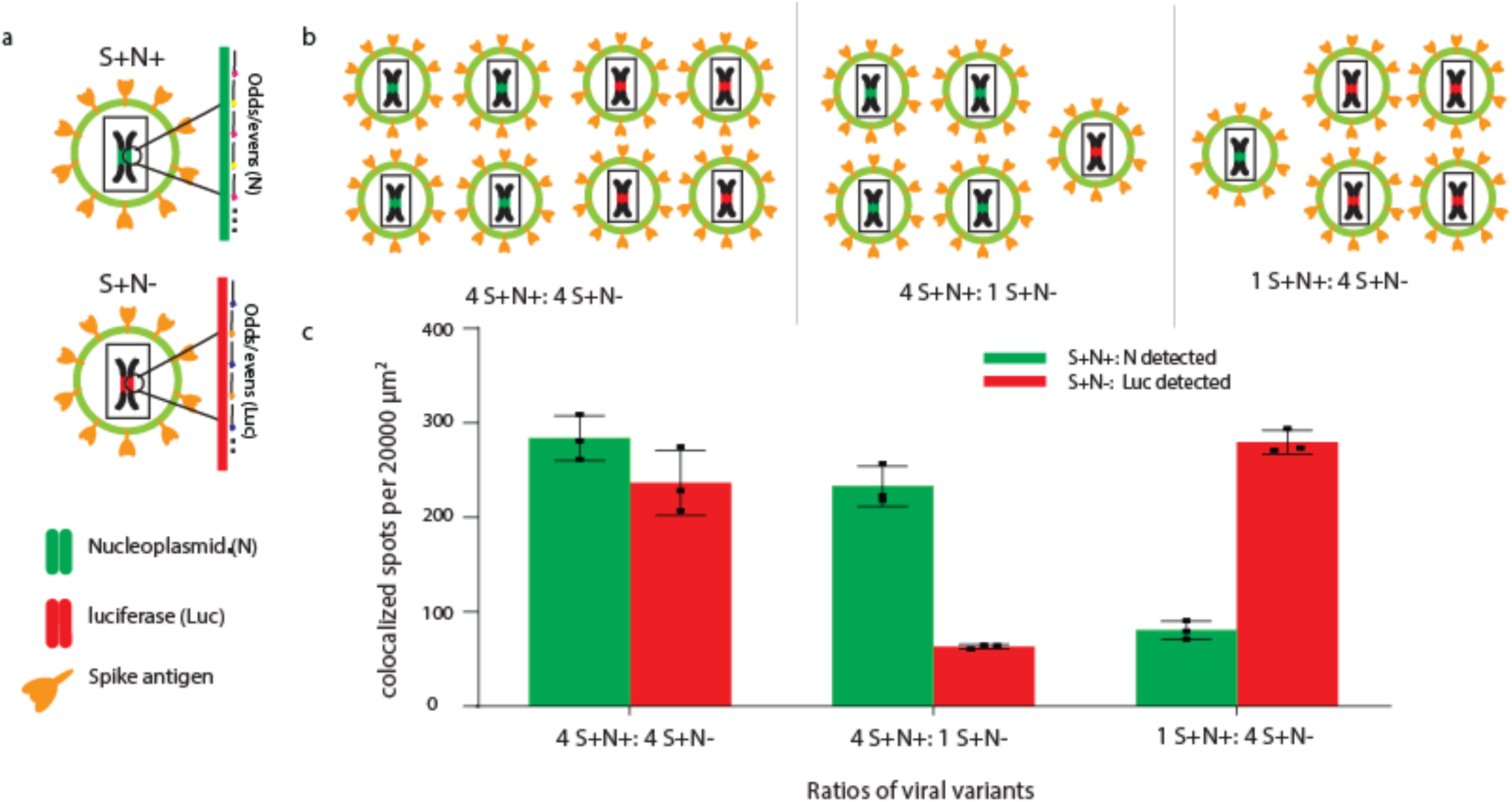
Multiplexed virus detection by TurboFISH. a. Multiple virus strains with different genomes were detected by N gene probes and luciferase probes with two sets of dyes. b. Different amount ratios of two virions were detected to show in c. whether the FISH spots ratio of two targets matches the concentration ratio of two virions added. 3 biological replicates, 5 images from each replicate (mean ± SD).

Three concentration ratios based on RT-qPCR of the two virion types were used to test our system: 4:4, 1:4, and 4:1 of S+N+:S+N-. For the 4:4 ratio, the ratio of colocalized spots of S+N+ to S+N- was 1.209:1. For the 4:1 ratio, the ratio of colocalized spots of S+N+ to S+N- was 3.679:1. For the 1:4 ratio, the ratio of colocalized spots of S+N+ to S+N- was 1: 3.498, consistent with the concentration ratio of the two virions we added, (**Figure 4c**) indicating that raptamer FISH can simultaneously detect multiple virion types.

### Functional titering of remaining supernatant from virions captured by aptamers and ACE2 receptors

Infectious titering relies on cell-based assays. Still accuracy and consistency of such methods heavily depend on cell condition^13^. Our system achieves a cell-free proxy for infectious titers. To test whether the infectious viral particles are indeed getting captured, we assessed the infectivity of the supernatant after incubating the virus on the ACE2 receptor and the 1C aptamer. The virion capture steps of aptamers and ACE2 were performed as described in Figure 2. The uncaptured virions in the supernatant were aspirated and used to transduce ACE2-expressing HEK293T cells (HEK293T-ACE2). After incubating HEK293T-ACE2 cells with the virus for two days, the percentage of infected cells was measured with flow cytometry via ZsGreen expression from the viral genome (**Figure 5a**). Wildtype HEK293FT cells were infected simultaneously to set up the gate of uninfected cells. Plates not coated with aptamers (uncoated) were used to do normalization and measure the total amount of infectious particles. The percentage of infected cells was used to quantify the infectivity of different samples. Random aptamers were used as a negative control.

**Figure 5.**
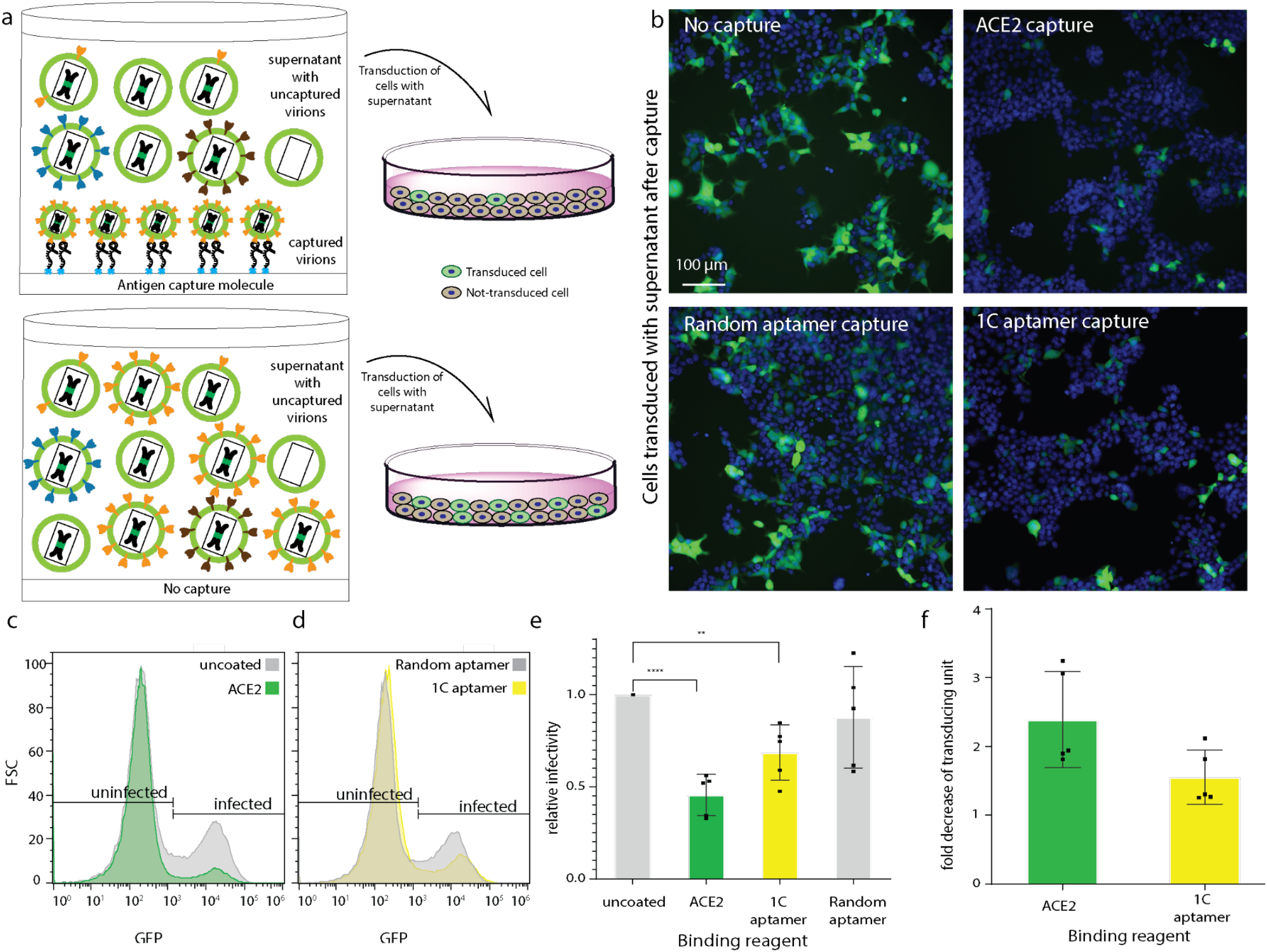
Functional titering of uncaptured virions from aptamers and ACE2 to prove the infectivity of captured virions. a. Uncaptured supernatant virions were aspirated out and used to infect cells. Flow cytometry was performed to count the percentage of infected cells. b. Images of cells infected by uncaptured virions from different binding reagents. The cell nucleus was stained with DAPI. c,d. Histogram plots of flow cytometry data, uninfected/infected cells were gated based on viral infection of 293FT cells. The ACE2 group was compared to the uncoated group, and the 1C aptamer group was compared to the random aptamer group. e. Normalization of uncaptured virions infectivity, percent of infected cells in different binding reagents was normalized to the uncoated group. f. Fold decrease of TU/mL of ACE2 and 1C aptamer groups were compared to the uncoated group. n = 5 biological replicates (mean ± SD). *p < 0.05, **p < 0.01, ***p < 0.001, ****p < 0.0001.

The percentage of infected cells decreased significantly after virion capture by ACE2 and 1C aptamers, compared to the infectivity of viruses that were not captured, suggesting that the infectious particles are being captured. Random aptamers had similar infectivity as the no capture group, which indicated that such capture is highly specific (**Figure 5b, Supplementary Figure 7**). Based on the flow cytometry data, two populations of uninfected/infected cells were gated in the histogram (**Figure 5c, 5d**). There was a 65.751% decrease in the percent of infected cells in the ACE2 group compared to the uncoated group (**Figure 5c**) and a 36.863% decrease in the percent of infected cells in the 1C aptamers group compared to the random aptamers group (**Figure 5d**). The random aptamer capture was comparable to the non-specific capture on uncoated plates. The percentage of infected cells in ACE2, 1C, and random aptamers were normalized to that of the uncoated group (**Figure 5e**). Percentages of infected cells were transformed to TU/mL using the viral quantification formula ^25^. A 2.392 fold reduction and a 1.552 fold reduction in TU/mL were observed in the ACE2 group and 1C aptamer group, respectively, compared to the uncoated group (**Figure 5f**).

## Discussion

Here, we outline raptamer FISH, a cell-free platform for the capture and direct detection of infectious virions using aptamers for specific capture of virions, and TurboFISH for rapid and specific detection of the viral genome. This system offers several advantages over current technologies: 1. The DNA aptamers are thermostable, inexpensive, and can reduce batch-to-batch variability. 2. TurboFISH allows for direct and specific detection of virions using standard epifluorescent microscopy. 3. Not all viruses are infectious; in this system, detection requires the presence of a surface antigen and genome, thus providing a cell-free proxy for infectious titer. 4. Finally, our image processing uses publicly available Matlab software, selecting for size (to exclude debris and virus aggregates), probe colocalization for specificity of detection, and use of multiple probes to discriminate different viruses in the sample.

We tested our technology on spike-pseudotyped lentiviruses bearing either a short region of the nucleocapsid gene from the SARS-CoV-2 genome or the luciferase gene. Our platform demonstrated specific spike-containing virus capture using ACE2 and 1C aptamer. Further, it can detect viral genomes in 15 minutes and discriminate between multiple genomes. Importantly, we demonstrated that ACE2 and 1C capture infectious virions, as viral supernatants after incubation with these agents relative to negative controls. These properties make the paired aptamer capture and FISH detection of virions a viable tool for the cell-free determination of infectious titer.

RT-PCR is most frequently used for viral measurements due to its sensitivity, which can detect as little as 100 copies of viral RNA per milliliter of transport media^29,30^. However, the measurement of genome-containing units fails to account for infectivity (i.e., a virus that contains a genome but the surface antigen is damaged or mutant and therefore cannot infect a cell^31^). The infectivity is important for diagnostics because a person may have non-infectious virions or unpackaged viral nucleic acids remaining in their body but still produce a positive test^9^. Likewise, the ELISA-based methods detect only the presence of surface antigen and do not inform on infectivity (i.e., virions may contain surface antigen without genome^32,33^). Alternatively, our capture and detection system, although theoretically single-molecule resolution, is limited by the capture efficiency of the aptamer or the antibody and, thus, will have lower sensitivity. However, this system combines the benefits of RT-PCR for genome detection and ELISA for surface antigen detection to inform on infectivity without the use of cells.

A major strength of this system is that aptamers may be selected via SELEX^34,35^ for any surface antigen, and RNA FISH probes may be designed to target different viral strains^20,27^. These properties will allow the system to be rapidly customized and applied to emerging pathogens and any desired viral vector without the need for fluorescent transgenes that are either not present in natural pathogens or not permitted in the minimum payload for gene therapies^36,37^.

In summary, we report the development of a direct, fast, and sensitive assay for virus detection and cell-free determination of infectious titers with no thermal requirement. This assay is unique because it first captures viruses bearing an intact coat protein using an aptamer, then detects genomes directly in individual virions using FISH–thus selecting for “infectious” particles. Importantly, we demonstrate that the virions captured are “infectious,” thus serving as a proxy of infectious titer. This measurement will circumvent the need for cell-based assays to determine infectious titers for diagnostic and biotechnology settings.

## Methods

### Tissue culture conditions

HEK293FT cells (ThermoFisher cat# R700-07) were handled according to the manufacturer’s protocol and maintained in high glucose DMEM with GlutaMax (FisherSci cat# 10566016), 10% FBS, 1% MEM Non-Essential Amino Acids (ThermoFisher cat# R70007), 1% supplementary GlutaMax (FisherSci cat# 35-050-061), 1% MEM sodium pyruvate (ThermoFisher cat# 11360070), 1% Pen-Strep (FisherSci cat# BW17-602E), and 500 ug/mL geneticin (ThermoFisher cat# 10131035). HEK293T-ACE2 cells described previously^25^ (BEI cat# NR-52511) were cultured in the HEK293FT media without geneticin. Cell lines tested PCR-negative for mycoplasma contamination (ATCC cat# 30-1012K).

### Plasmids

The following plasmids were ordered as part of a lentiviral kit (BEI cat# NR-52948) described previously^25^: Spike (BEI cat# NR-52514), Luc2-ZsGreen (BEI cat# NR-52516), Hgpm2 (BEI cat# NR-52517), Tat1b (BEI cat# NR-52518), and Rev1b (BEI cat# NR-52519). VSV-G plasmid (Addgene cat# 12259) was a gift from the lab of Jiahe Li. To prepare the Spike G614 Δ19 plasmid, we generated 2 PCR amplicons using mutagenic primers, Q5 polymerase (NEB cat# M0491) and Spike plasmid (BEI cat# NR-52514) as a template. PCR reactions were digested in DpnI, and the amplicons were joined via Gibson assembly (NEB cat# E2611). The coding sequences of generated plasmids were sequence-verified. To prepare the N gene inserted plasmids (CoV2Ngene-Luc2-ZsGreen) to model SARS-COV-2, we removed an 1131 bp segment of luciferase in Luc2-ZsGreen (BEI cat# NR-52516), and the N gene (1260 bp, IDT 2019-nCoV_N_Positive Control Plasmid cat#10006625) was amplified by mutagenic primers. The remaining part of Luc2-ZsGreen (BEI cat# NR-52516) and the amplified N gene were ligated via Gibson assembly (NEB cat# E2611). The coding sequences of generated plasmids were sequence-verified.

### Generating Pseudotyped Lentiviral Particles

The protocol follows the previously described in literature^25^ with several modifications. HEK293FT cells were seeded in media without geneticin at a density such that cells were 50% confluent the next day. 16 to 24 hours after seeding, cells were transfected with the lentiviral plasmids using BioT (Bioland cat# B01-01) according to the manufacturer’s protocol. The number of plasmids per transfection was at a 1:2/1:9/1:6 mass ratio for the lentiviral backbone (Luc2-ZsGreen or N-ZsGreen), helper plasmids (Hgpm2, Tat1b, and Rev1b), and viral entry protein (Spike G614 Δ19, or VSV-G), respectively. Media was changed 18 hours post-transfection (hpt). Viral supernatant was collected at 60 hpt and kept and centrifuged at 500 x g for 10 min to clear cell debris. Cleared supernatant was immediately frozen at -80 °C in aliquots.

### Viral quantification

Viruses were quantified via a previously described RT-qPCR method^38^, where we only deviated by using different DNase I (ThermoFisher #AM1907) digestion, reverse transcriptase (FisherSci #18-080-044), and plasmid (Luc2-ZsGreen) for generating standard curves. A CFX96™ Optics Module, C1000 Touch™ Thermal Cycler (BIO-RAD) was used. Finally, viruses were functionally titered via HEK293T-ACE2 infection as previously described^25^, with the only deviation being that the HEK293T-ACE2 cells were cultured in HEK293FT media without geneticin. 48h post-infection, cells were analyzed with the Attune NxT flow cytometer for fluorescence, indicating a successful viral infection. The transducing units of infectious virions were obtained from the P (percent of infective positive cells) based on the formula:

TU/mL = -ln(1 - [P/100]) * (# cells in well/volume of viral stock added to well in mL),

### Chemical immobilization of biotin to a chambered coverglass

To generate the biotinylated coverglass necessary to run and image raptamer FISH, an 8-well chambered coverglass (ThermoFisher #155409) was conditioned in an oxygen-plasma cleaner to introduce the active oxygen group to the slide. The slide was cleaned with water and then 100% isopropanol before the reaction. The slide was treated by oxygen at 300 mTorr pressure for 1 min and then by plasma power at 50 mW for 1 min. After the oxygen plasma, silane-PEG-biotin (10 mg/mL) dissolved in 99% ethanol was added to react with the oxygen group and incubated for 4-6 h at RT. The slide was washed with 99% ethanol three times after silane-PEG-biotin treatment. The dried slide was kept at 4 °C overnight for use the next day.

### ELISA assay

Recombinant spike protein (BPS bioscience #100810) was captured by ACE2 receptor protein (Acro biosystems #AC2-H82F9) and aptamers (CoV2-RBD-1C and CoV-RBD-4C) targeting receptor-binding domains of SARS-CoV-2 spike glycoprotein which were discovered in Song *et al*.^22^ Biotin-modified aptamers and ACE2 were bound to streptavidin-coated polystyrene plates (ThermoFisher #PI15500). The aptamers and ACE2 captured the recombinant spike protein, and then SARS-CoV-2 Spike Protein Monoclonal Antibody [H6] HRP-conjugated (AssayGenie #PACOV0004) was used to detect the captured spike protein. The ELISA procedure to detect spike protein was conducted according to the streptavidin-coated plate manufacturer’s protocol with the following changes: Aptamers (10 uM), spike protein (1 ug/mL), and monoclonal antibodies (1 ug/mL) were dissolved in the binding buffer (1X PBS, 0.55 mM MgCl_2_). ACE2 protein (500 ng/mL) was dissolved in the wash buffer (25mM Tris, 150 mM NaCl, 0.1% FBS, 0.05% Tween-20). The wells were pre-washed with 1X PBS for 10 minutes and then the binding buffer for 10 minutes before adding the aptamers. Aptamers were heated at 90 °C for 10 minutes and placed on ice for 10 minutes. The aptamers and ACE2 protein had 2 hours of binding time. Both the spike protein and the antibody had 30 minutes of binding time. The plates were pre-incubated with HEK293T-ACE2 cell media with 0.55 mM MgCl_2_ for 10 min. The virus was prepared in HEK293T-ACE2 cell media with 0.55 mM MgCl_2_ and ssDNA. The virus was added to the plates and incubated for 45 min, and the plates were washed 3 times with binding buffer after virus incubation. For biotinylated coverglass, streptavidin protein (100 μg/mL) was added first and incubated for 1 h, and the remaining steps were the same as described above.

### RNA extraction, RT-PCR and RT-qPCR

Methods were adapted from Zhang *et al^38^*. Frozen viral aliquots were thawed and treated with RNase A (EN0531 ThermoFisher, diluted 10X) for 1 h at 37 °C to degrade RNAs that were not packaged inside the virion. RNA was extracted using Trizol (ThermoFisher #15596026) and resuspended in NF water. DNase (ThermoFisher #AM1907) was used to remove the contaminant DNA. RNA was reverse transcribed (FisherSci #18-080-044). The Luna qPCR (NEB #M3004) was used to quantify the generated cDNA. OneTaq RT-PCR mix (NEB #E5315) was used to amplify the generated cDNA.

### Raptamer FISH

The steps after immobilization were performed as described previously in the literature^16^. The coverglass was washed 3 times with the binding buffer after virions capture. To fix virus samples, -20 °C methanol was added to samples, which were then incubated at -20 °C for 10 min. 500 μM of probe mixture was mixed with the hybridization buffer (10% dextran sulfate/10% formamide/2X SSC) and incubated with the fixed samples for 5 min. The samples were washed 2 times with the wash buffer (2X SSC, 10% formamide) then 2X SSC was added for subsequent imaging.

### Image acquisition

Microscopy was performed using a Nikon inverted research microscope eclipse Ti2-E/Ti2-E/B using a Plan Apo λ 20X/0.75 objective or Plan Apo λ 100X/1.45 oil objective.The Epi-fi LED illuminator linked to the microscope assured illumination and controlled the respective brightness of four types of LEDs of different wavelengths. Images were acquired using the Neutral Density (ND16) filter for probes coupled with Alexa 488, Alexa 594, Alexa 647, cy3. Images were acquired and processed using ImageJ. Images acquired using the Neutral Density (ND16) filter are false-colored gray. Nucleus of live cells were stained by NucBlue™ Live Cell Stain ReadyPorbes™ reagent (ThermoFisher #R38605).

### Image analysis and quantification

After imaging, we put our data through an image analysis pipeline for semi-automated spot recognition. The pipeline, developed in Matlab, can be divided into the three main steps of binarization, colocalization and size thresholding. Briefly, in the first binarization step we manually determine a threshold for each fluorescence channel so the number of spots can be localized well in positive control. Images of each channel share the same thresholding level, which is applied to the images to binarize them. The binarized images are then overlapped and intersected in order to find the common spots to the two channels (**Figure 3 and Figure 4**). Eventually, in the last step we set a cut-off level for the particle area, in order to exclude viral aggregates from the analysis and limit the colocalized group to just individual virions (**Supplementary Figure 7**).

## Author Contributions

Conceived of and designed the experiments: YL, JLP, DB, and SHR. Performed the experiments: YL, JLP, DB, and KN. Analyzed the data: YL, JLP, DB, KN, CAM, OF. Wrote the paper: YL and SHR with contributions from all authors.

## Acknowledgements

The authors would like to acknowledge support from the National Science Foundation (NSF) RAPID Program grant (NSF-2032533) as well as the NSF Partnerships for Innovation program (NSF-2141135).

