## Supporting information for "Paired aptamer capture and FISH detection of individual virions enables cell-free determination of infectious titer"

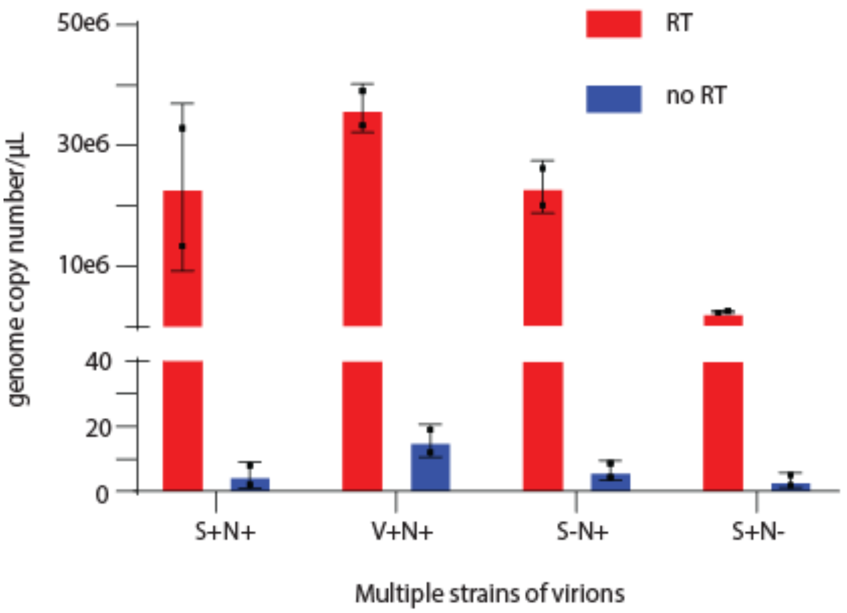

**Supplementary Figure 1:** RT-qPCR quantification of lentiviral vectors used in this study. Multiple virus strains were quantified after production, genome copies number/ µL were shown, no RT means no reverse transcription was performed, S+ represents presence of spike protein on the virus surface, N+ represents presence the Nucleocapsid gene in the lentiviral genome, S- represents absence of any viral coat protein, and V represents presence of VSVG on the virus surface. n = 2 biological replicates (bars represent mean ± SD).

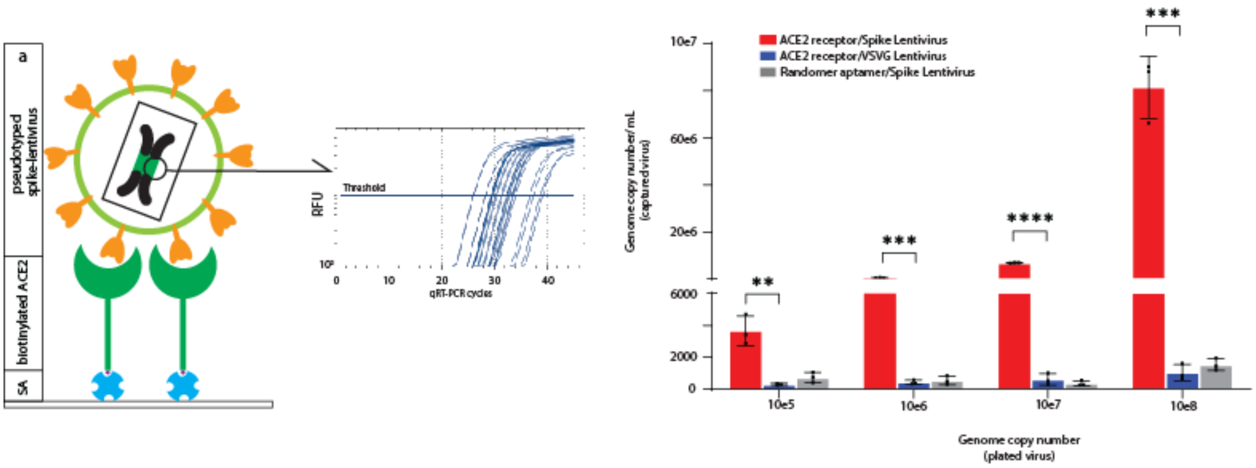

**Supplementary Figure 2:** RT-qPCR for capture efficiencies of SARS-COV-2 spike pseudotyped lentivirus by ACE2 receptor. a. ACE2 receptors were conjugated to a streptavidin-coated polystyrene plate. Following lentivirus were captured by the receptors, RNA was extracted, and RT-qPCR was performed to measure the amount of captured virions. b. Bar graph of genome copies number per mL of spike and VSVG virions captured by ACE2 receptors and random aptamers. n = 3 biological replicates (mean ± SD). *p < 0.05, **p < 0.01, ***p < 0.001, ****p < 0.0001.

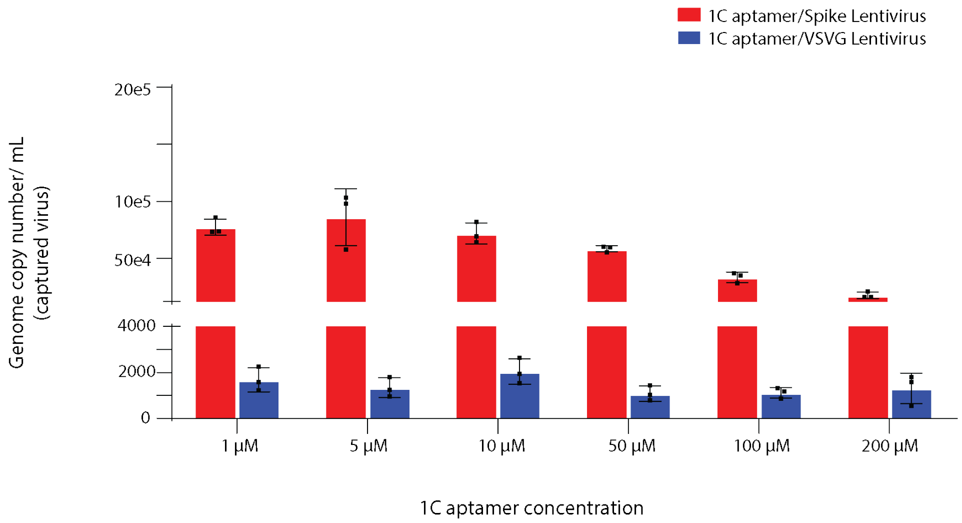

**Supplementary Figure 3:** Saturation test of 1C aptamers concentration. Six different concentrations of 1C aptamers ranging from 1 μM to 200 μM were tested, VSVG lentivirus was used as a negative control to indicate nonspecific binding to 1C aptamer, the genome copy number per mL of spike and VSVG virions captured by difference concentration of 1C aptamers were shown in the bar graph. n=3 biological replicates (mean ± SD).

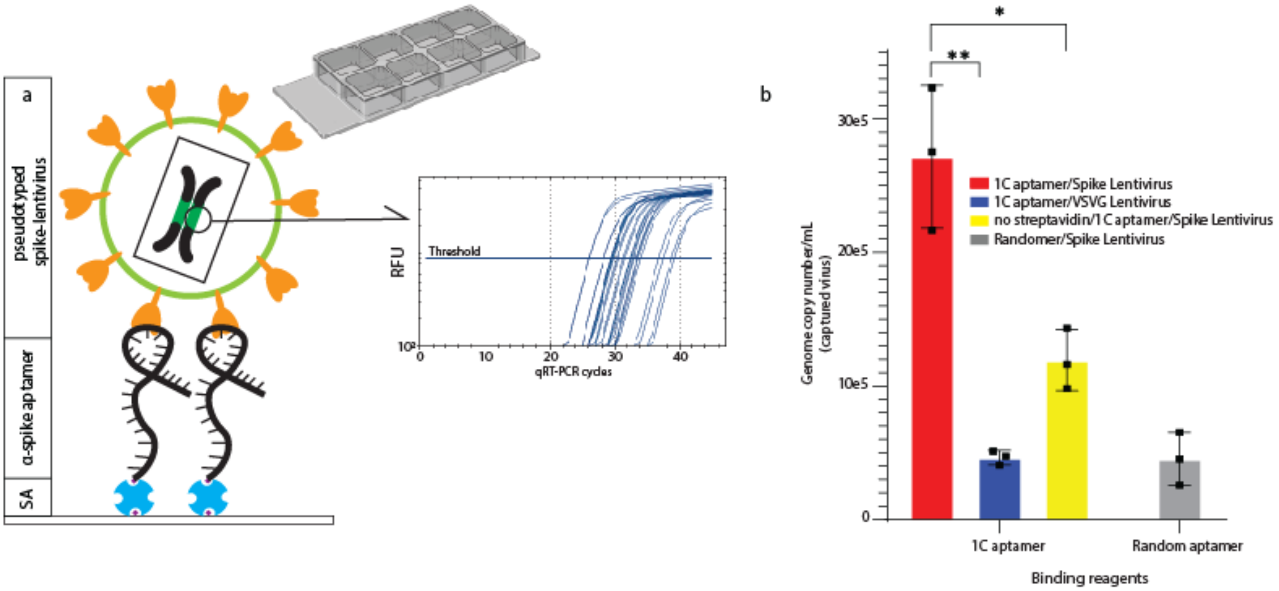

**Supplementary Figure 4:** RT-qPCR for capture efficiencies of SARS-CoV-2 spike pseudotyped lentivirus by 1C aptamers on chambered coverglass. To prove virions can be captured on coverglass applicable for FISH, chambered coverglass was biotinylated by oxygen plasma, then streptavidin was added, and biotinylated 1C aptamer was immobilized. The spike-pseudotyped lentivirus and the VSVG-pseudotyped lentivirus were added to the coverglass at concentrations of $10^6$ genome copies per µl. After washing, we extracted the RNA from captured virions and performed RT-qPCR targeting the CMV promoter (common between both lentivirus vectors) to determine capture efficiency. No streptavidin-treated sample was used as the negative control. The genome copy number per mL of spike and VSVG virions captured by 1C aptamers and random aptamers were shown in the bar graph. n=3 biological replicates (mean ± SD). *$p < 0.05$, **$p < 0.01$, ***$p < 0.001$, ****$p < 0.0001$.

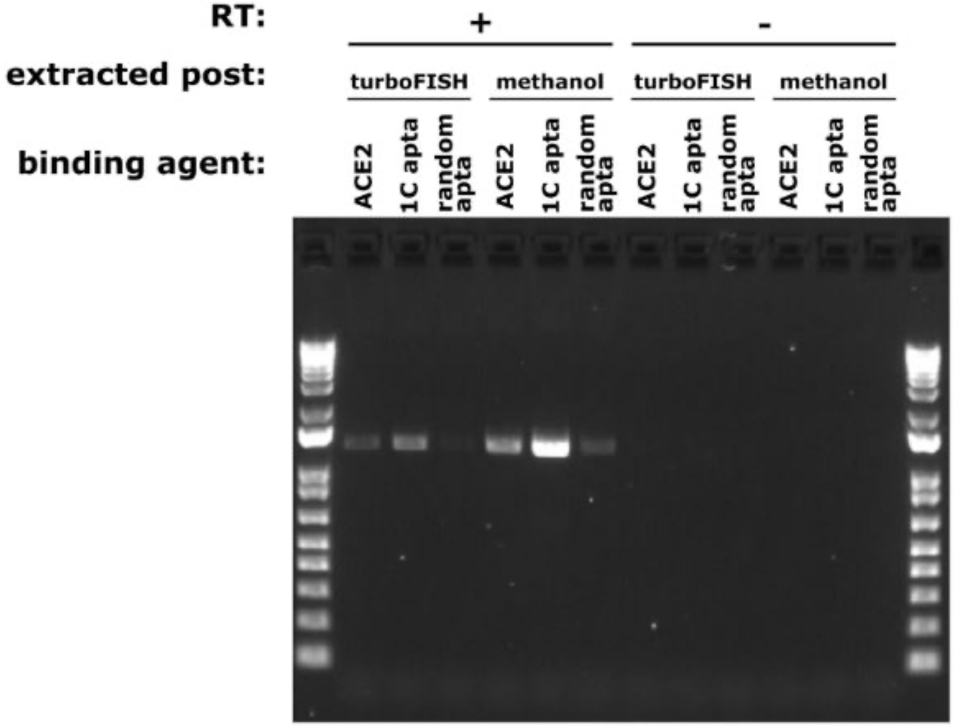

**Supplementary Figure 5:** DNA agarose gel electrophoresis to prove the integrity of viral genome post-methanol fixation and post-TurboFISH treatment. RT+ means reverse transcription was performed. Virions were captured, and viral mRNA was extracted post methanol or TurboFISH, RT-PCR targeting the N gene of S+N+ was performed to detect the 1260 bp band in different binding reagents.

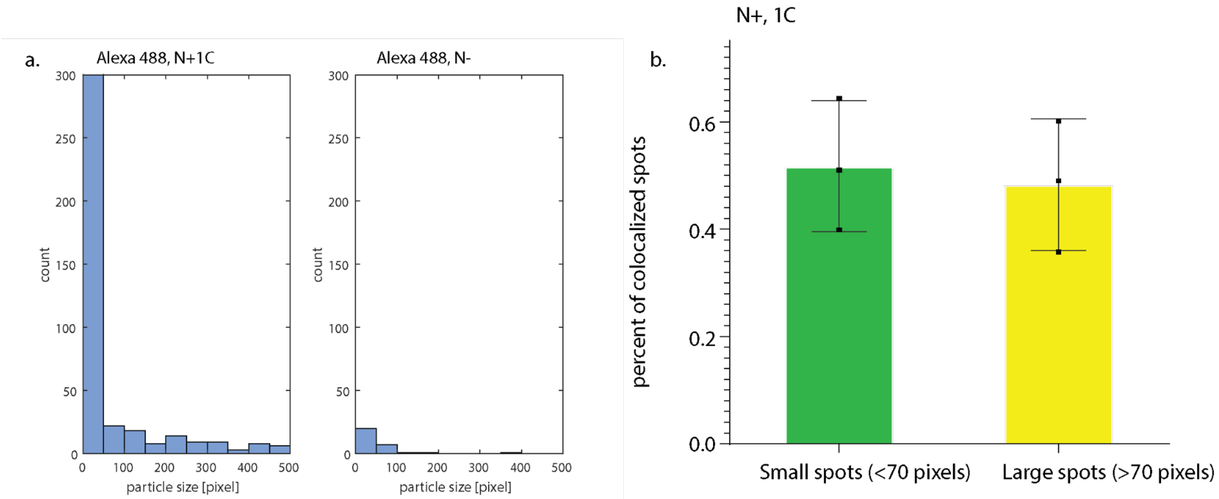

**Supplementary Figure 6:** a. Particle size (pixel) distribution of FISH spots in Figure 3. After the binarization step, the particles detected in the alexa 488 channel were measured in Matlab, histogram of size distribution was shown. 1C aptamer group capture spike coated lentivirus with N gene was compared to no 1C aptamer group capture spike coated lentivirus without N gene. b. Percent of colocalized spots for small spots (<70 pixels) and large spots (>70 pixels) in 1C captured spike coated lentivirus with N gene. n=3 biological replicates (mean ± SD).

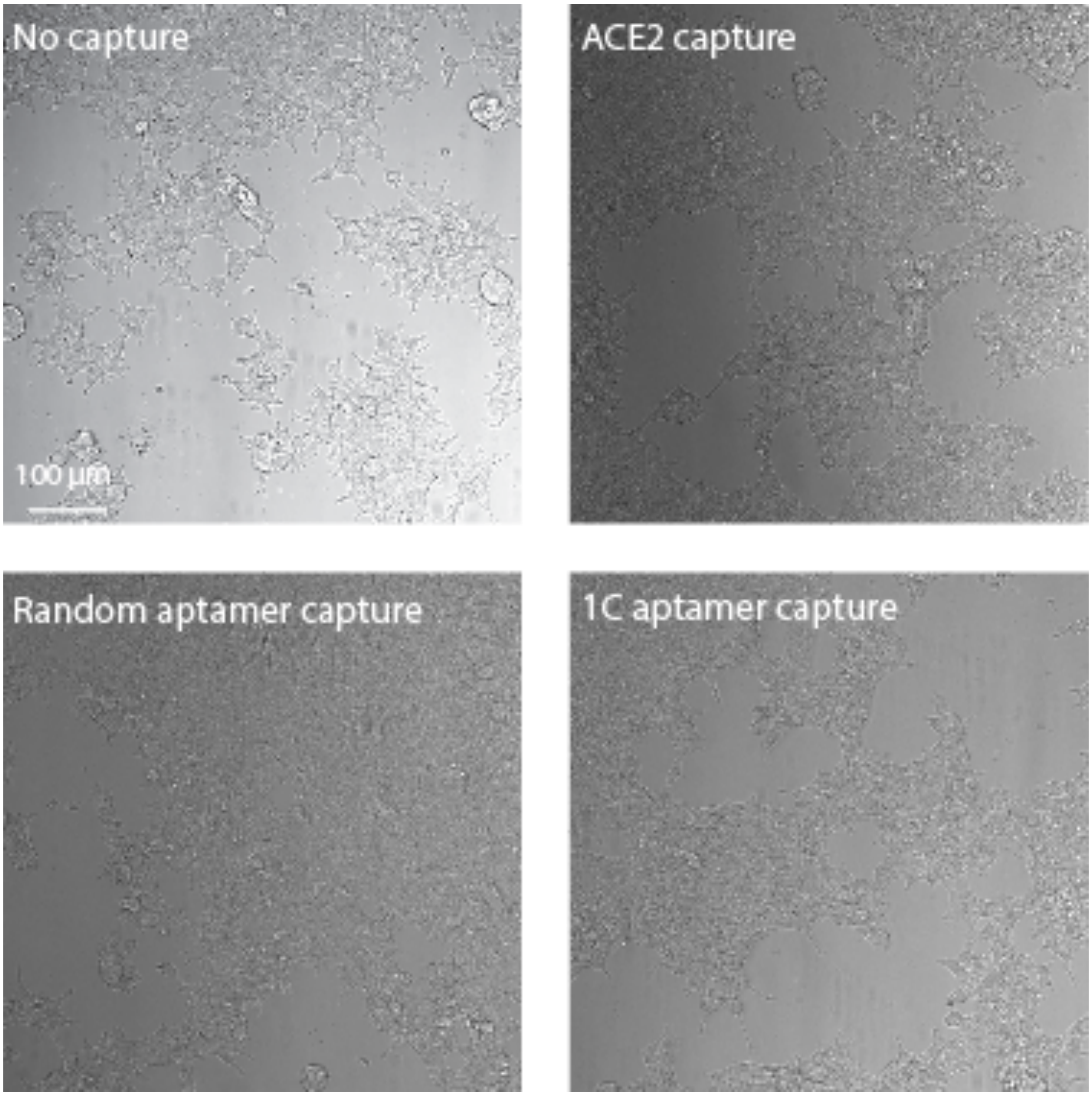

**Supplementary Figure 7:** Brightfield images of monolayer cells in Figure 5b.

**Supplementary Methods**

Table 1. Primers and probes used for RT-qPCR to quantify captured virions

| Name | Amplicon length(bp) | Description | Sequence 5' to 3' |
|---|---|---|---|
| CMV promoter | 104 bp | Forward | TCACGGGGATTTCCAAGTCTC |
| | | Reverse | AATGGGGCGGAGTTGTTACGAC |
| | | Probe | /56-FAM/AAACAAACT/ZEN/CCCATTGACGTCA/3IaBkFQ/ |

Table 2. Primers used for RT-PCR to amplify N gene

| Name | Amplicon length(bp) | Description | Sequence 5' to 3' |
|---|---|---|---|
| Nucleoplasmids | 1260 bp | Forward | ATGTCTGATAATGGACCCCA |
| | | Reverse | TTAGGCCTGAGTTGAGTC |

**Two-step RT-qPCR**

Use SuperScript III Reverse Transcriptase (FisherSci #18-080-044), RNaseOUT (FischerSci #10777-019) and Luna qPCR (NEB #M3004). Reverse transcription was performed by mixing 7.5 µl DNase digestion product (10 pg-500 ng mRNA), 1µl 10mM dNTPs, 1µl 2 µM CMV-reverse primer, and Nuclease free water up to 13µl, followed by incubation at 65°C for 3min and ice for 1 min. The sample was placed on ice and a mix containing 1 µl 0.1M DTT, 4µl 5x First-Strand Buffer, 1µl RNaseOUT, 1ul SuperScript III RT (200 U/ul) (replace with Nuclease free water for RT negative control) and water up to 7µl was added. The samples were then incubated at 50°C for 60min followed by 70°C for 15min. Amplification of 1µl cDNA (RT mix) using primers and probes described in Table 1 was performed using in a 20µl reaction containing 10µl Master Mix, 0.8 ul forward, 0.8 ul reverse 10 uM CMV primers, 0.4 ul 10 uM CMV probe and water up to 20µl. The thermal cycling steps were: 95°C for 1min, and 45 cycles of 95°C for 15s and 60°C for 30s. qPCR was performed on a CFX96™ Optics Module, C1000 Touch™ Thermal Cycler (BIO-RAD) using the CFX Manager™ Software v3.1.

**One-step RT-PCR**

For reverse transcription and PCR we used the OneTaq RT-PCR mix (NEB #E5315) according to the manufacturer's instructions. Reactions of 25µl were formed by mixing DNA digestion product (mRNA up to 1µg), One*Taq* One-Step Reaction Mix (2X) 12.5 µl, One*Taq* One-Step Enzyme Mix (25X) 1 µl, 1 µl 10 µM forward primer and reverse primer listed in Table 2 and Nuclease free water up to 25µl. The thermal cycling steps were: 48°C for 2min, 94°C for 1min and 30 cycles of 94°C for 15 s and 50°C for 30s, 68°C for 75s. After cycles final extension was 68°C for 5 min and hold at 10 °C. RT-PCR was performed on a T100 Thermal Cycler machine (BIO-RAD).

**Supplementary Note: Sequences**
All the plasmids used to generate lentivirus particles are listed below, all the modified sequences were sequence-verified.

Vector name: Hgpm2

agcttggcccattgcatacgttgtatccatatcataatatgtacatttatattggctcatgtccaacattaccgccatgttgacattgatt
attgactagttattaatagtaatcaattacggggtcattagttcatagcccatatatggagttccgcgttacataacttacggtaaatgg
cccgcctggctgaccgcccaacgacccccgcccattgacgtcaataatgacgtatgttcccatagtaacgccaatagggactttc
cattgacgtcaatgggtggagtatttacggtaaactgcccacttggcagtacatcaagtgtatcatatgccaagtacgccccctatt
gacgtcaatgacggtaaatggcccgcctggcattatgcccagtacatgaccttatgggactttcctacttggcagtacatctacgta
ttagtcatcgctattaccatggtgatgcggttttggcagtacatcaatgggcgtggatagcggtttgactcacggggatttccaagtct
ccaccccattgacgtcaatgggagtttgttttggcaccaaaatcaacgggactttccaaaatgtcgtaacaactccgccccattga
cgcaaatgggcggtaggcgtgtacggtgggaggtctatataagcagagctcgtttagtgaaccgtcagatcgcctggagacgcc
atccacgctgttttgacctccatagaagacaccgggaccgatccagcctcccctcgaagctgatcctgagaacttcagggtgagt
ctatgggacccttgatgttttctttccccttcttttctatggttaagttcatgtcataggaaggggagaagtaacagggtacacatattga
ccaaatcagggtaattttgcatttgtaattttaaaaaatgctttcttcttttaatatacttttttgtttatcttatttctaatactttccctaatctct
ttctttcagggcaataatgatacaatgtatcatgcctctttgcaccattctaaagaataacagtgataatttctgggttaaggcaatagc
aatatttctgcatataaatatttctgcatataaattgtaactgatgtaagaggtttcatattgctaatagcagctacaatccagctaccatt
ctgcttttattttatggttgggataaggctggattattctgagtccaagctaggcccttttgctaatcatgttcatacctcttatcttcctcc
cacagctcctgggcaacgtgctggtctgtgtgctggcccatcactttggcaaagaattctagactgccatgggcgcccgcgcctc
cgtgctgtccggcggcgagctggacaagtgggagaagatccgcctgcgccccggcggcaagaagcagtacaagctgaagc
acatcgtgtgggcctcccgcgagctggagcgcttcgccgtgaaccccggcctgctggagacctccgagggctgccgccagat
cctgggccagctgcagccctccctgcaaaccggctccgaggagctgcgctccctgtacaacaccatcgccgtgctgtactgcg
tgcaccagcgcatcgacgtgaaggacaccaaggaggccctggacaagatcgaggaggagcagaacaagtccaagaagaa
ggcccagcaggccgccgccgacaccggcaacaactcccaggtgtcccagaactaccccatcgtgcagaacctgcagggcc
agatggtgcaccaggccatctcccccgcaccctgaacgcctgggtgaaggtggtggaggagaaggccttctcccccgaagt
catccccatgttctccgccctgtccgagggcgccacccccaggacctgaacaccatgctgaacaccgtgggcggccaccag
gccgccatgcagatgctgaaggagaccatcaacgaggaggccgccgagtgggaccgcctgcaccccgtgcacgccggccc
catcgcccccggccagatgcgcgagccccgcggctccgacatcgccggcaccacctccaccctgcaagagcagatcggctg
gatgacccacaacccccccatccccgtgggcgagatctacaagcgctggatcatcctgggcctgaacaagatcgtgcgcatgt
actcccccacctccatcctggacatccgccagggccccaaggagcccttccgcgactacgtggaccgcttctacaagaccctg
cgcgccgagcaggcctcccaggaggtaaagaactggatgaccgagaccctgctggtgcagaacgccaaccccgactgcaa
gaccatcctgaaggccctgggccccggcgccaccctggaggagatgatgaccgcctgccagggcgtgggcggccccggcc
acaaggcccgcgtgctggccgaggccatgtcccaagtcaccaaccccgccaccatcatgatccagaagggcaacttccgcaa
ccagcgcaagaccgtgaagtgcttcaactgcggcaaggagggccacatcgccaagaactgccgcgcccccgcaagaagg
gctgctggaagtgcggcaaggagggccaccagatgaaagattgtactgagagacaggctaatttttttagggaagatctggccttc
ccacaagggaaggccagggaattttcttcagagcagaccagagccaacagccccaccagaagagagcttcaggtttggggaa
gagacaacaactccctctcagaagcaggagccgatagacaaggaactgtatcctttagcttccctcagatcactctttggcagcg
acccctcgtcacaataaagatcggtggccagctgaaggaggccctgctggacaccggcgccgacgacaccgtgctggagga
gatgaacctgcccggccgctggaagcccaagatgatcggcggcatcggcggcttcatcaaagtccgccagtacgaccagatc
ctgatcgagatctgcggccacaaggccatcggcaccgtgctggtgggccccacccccgtgaacatcatcggccgcaacctgct
gacccagatcggctgcaccctgaacttccccatctcccccatcgagaccgtgcccgtgaagctgaagcccggcatggacggc
cccaaagtcaagcagtggcccctgaccgaggagaagatcaaggccctggtggagatctgcaccgagatggagaaggaggg
caagatctccaagatcggccccgagaacccctacaacacccccgtgttcgccatcaagaagaaggactccaccaagtggcgc
aagctggtggacttccgcgagctgaacaagcgcacccaggacttctgggaggtgcagctgggcatcccccaccccgccggc
ctgaagcagaagaagtccgtgaccgtgctggacgtgggcgacgcctacttctccgtgcccctggacaaggacttccgcaagta
caccgccttcaccatcccctccatcaacaacgagacccccggcatccgctaccagtacaacgtgctgccccagggctggaag

ggctcccccgccatcttccagtgctccatgaccaagatcctggagcccttccgcaagcagaaccccgacatcgtgatctaccag
tacatggacgacctgtacgtgggctccgacctggagatcggccagcaccgcaccaagatcgaggagctgcgccagcacctgc
tgcgctggggcttcaccacccccgacaagaagcaccagaaggagcccccccttcctgtggatgggctacgagctgcaccccga
caagtggaccgtgcagcccatcgtgctgcccgagaaggactcctggaccgtgaacgacatccagaagctggtgggcaagctg
aactgggcctcccagatctacgccggcatcaaagtccgccagctgtgcaagctgctgcgcggcaccaaggccctgaccgagg
tggtgcccctgaccgaggaggccgagctggagctggccgagaaccgcgagatcctgaaggagcccgtgcacggcgtgtact
acgacccctccaaggacctgatcgccgagatccagaagcagggccagggccagtggacctaccagatctaccaggagccct
tcaagaacctgaagaccggcaaatacgcccgcatgaagggcgcccacaccaacgacgtgaagcagctgaccgaggccgtg
cagaagatcgccaccgagtccatcgtgatctggggcaagactcccaagttcaagctgcccatccagaaggagacctgggagg
cctggtggaccgagtactggcaggccacctggatccccgagtgggagttcgtgaacacccccccccctggtgaagctgtggtac
cagctggagaaggagcccatcatcggcgccgagaccttctacgtggacggcgccgccaaccgcgagaccaagctgggcaa
ggccggctacgtgaccgaccgcggccgccagaaggtggtgcccctgaccgacaccaccaaccagaagaccgagctgcag
gccatccacctggccctgcaagactccggcctggaggtgaacatcgtgaccgactcccagtatgcattgggcatcatccaggc
ccagcccgacaagtccgagtccgagctggtgtcccagatcatcgagcagctgatcaagaaggagaaggtgtacctggcctgg
gtgcccgcccacaagggcatcggcggcaacgagcaggtggacaagctggtgtccgccggcatccgcaaggtgctgttcctgg
acggcatcgacaaggcccaggaggagcacgagaagtaccactccaactggcgcgccatggcctccgacttcaacctgcccc
ccgtggtggccaaggagatcgtggcctcctgcgacaagtgccagctgaagggcgaggccatgcacggccaggtggactgctc
cccggcatctggcagctggactgcacccacctggagggcaaggtgatcctggtggccgtgcacgtggcctccggctacatcg
aggccgaggtgatccccgccgagaccggccaggagaccgcctacttcctgctgaagctggccggccgctggcccgtgaaga
ccgtgcacaccgacaacggctccaacttcacctccaccaccgtgaaggccgcctgctggtgggccggcatcaagcaggagtt
cggcatcccctacaacccccagtcccagggcgtgatcgagtccatgaacaaggagctgaagaagatcatcggccaagtccgc
gaccaggccgagcacctgaagaccgccgtgcagatggccgtgttcatccacaacttcaagcgcaagggcggcatcggcggc
tactccgccggcgagcgcatcgtggacatcatcgccaccgacatccagaccaaggagctgcagaagcagatcaccaagatc
cagaacttccgcgtgtactaccgcgactcccgcgaccccgtgtggaagggccccgccaagctgctgtggaagggcgagggc
gccgtggtgatccaggacaactccgacatcaaggtggtgccccgccgcaaggccaagatcatccgcgactacggcaagcag
atggccggcgacgactgcgtggcctcccgccaggacgaggactaacacatggaaaagattagtaaaacaccataggccgctc
tagaggatccaagcttatcgataccgtcgacctcgagggcccagatctaattcaccccaccagtgcaggctgcctatcagaaag
tggtggctggtgtggctaatgccctggcccacaagtatcactaagctcgctttcttgctgtccaatttctattaaaggttcctttgttccc
taagtccaactactaaactgggggatattatgaagggccttgagcatctggattctgcctaataaaaaacatttattttcattgcaatg
atgtatttaaattatttctgaatattttactaaaaagggaatgtgggaggtcagtgcatttaaaacataaagaaatgaagagctagttc
aaaccttgggaaaatacactatatcttaaactccatgaaagaaggtgaggctgcaaacagctaatgcacattggcaacagcccct
gatgcctatgccttattcatccctcagaaaaggattcaagtagaggcttgatttggaggttaaagttttgctatgctgtattttacattac
ttattgttttagctgtcctcatgaatgtcttttcactacccatttgcttatcctgcatctctcagccttgactccactcagttctcttgcttag
agataccacctttcccctgaagtgttccttccatgttttacggcgagatggtttctcctcgcctggccactcagccttagttgtctctgtt
gtcttatagaggtctacttgaagaaggaaaaacaggggggcatggtttgactgtcctgtgagcccttcttccctgcctcccccactca
cagtgacccggaatccctcgacatggcagtctagatcattcttgaagacgaaagggcctcgtgatacgcctatttttataggttaat
gtcatgataataatggtttcttagacgtcaggtggcacttttcggggaaatgtgcgcggaacccctatttgtttatttttctaaatacatt
caaatatgtatccgctcatgagacaataaccctgataaatgcttcaataatattgaaaaaggaagagtatgagtattcaacatttccg
tgtcgcccttattcccttttttgcggcattttgccttcctgtttttgctcacccagaaacgctggtgaaagtaaaagatgctgaagatca
gttgggtgcacgagtgggttacatcgaactggatctcaacagcggtaagatccttgagagttttcgccccgaagaacgttttccaat
gatgagcacttttaaagttctgctatgtggcgcggtattatcccgtattgacgccgggcaagagcaactcggtcgccgcatacact
attctcagaatgacttggttgagtactcaccagtcacagaaaagcatcttacggatggcatgacagtaagagaattatgcagtgct
gccataaccatgagtgataacactgcggccaacttacttctgacaacgatcggaggaccgaaggagctaaccgcttttttgcaca
acatgggggatcatgtaactcgccttgatcgttgggaaccggagctgaatgaagccataccaaacgacgagcgtgacaccacg
atgcctgtagcaatggcaacaacgttgcgcaaactattaactggcgaactacttactctagcttcccggcaacaattaatagactg
gatggaggcggataaagttgcaggaccacttctgcgctcggcccttccggctggctggtttattgctgataaatctggagccggtg
agcgtgggtctcgcggtatcattgcagcactggggccagatggtaagccctcccgtatcgtagttatctacacgacggggagtca
ggcaactatggatgaacgaaatagacagatcgctgagataggtgcctcactgattaagcattggtaactgtcagaccaagtttact
catatatactttagattgatttaaaacttcatttttaatttaaaaggatctaggtgaagatcctttttgataatctcatgaccaaaatccctt

aacgtgagttttcgttccactgagcgtcagaccccgtagaaaagatcaaaggatcttcttgagatcctttttttctgcgcgtaatctgc
tgcttgcaaacaaaaaaaccaccgctaccagcggtggtttgtttgccggatcaagagctaccaactctttttccgaaggtaactgg
cttcagcagagcgcagataccaaatactgttcttctagtgtagccgtagttaggccaccacttcaagaactctgtagcaccgccta
catacctcgctctgctaatcctgttaccagtggctgctgccagtggcgataagtcgtgtcttaccgggttggactcaagacgatagt
taccggataaggcgcagcggtcgggctgaacggggggttcgtgcacacagcccagcttggagcgaacgacctacaccgaac
tgagatacctacagcgtgagctatgagaaagcgccacgcttcccgaagggagaaaggcggacaggtatccggtaagcggca
gggtcggaacaggagagcgcacgagggagcttccagggggaaacgcctggtatctttatagtcctgtcgggtttcgccacctct
gacttgagcgtcgatttttgtgatgctcgtcaggggggcggagcctatggaaaaacgccagcaacggagatgcgccgcgtgcg
gctgctggagatggcggacgcgatggatatgttctgccaagggttggtttgcgcattcacagttctccgcaagaattgattggctcc
aattcttggagtggtgaatccgttagcgaggtgccgccggcttccattcaggtcgaggtggcccggctccatgcaccgcgacgc
aacgcggggaggcagacaaggtatagggcggcgcctacaatccatgccaacccgttccatgtgctcgccgaggcggcataa
atcgccgtgacgatcagcggtccaatgatcgaagttaggctggtaagagccgcgagcgatccttgaagctgtccctgatggtcgt
catctacctgcctggacagcatggcctgcaacgcgggcatcccgatgccgccggaagcgagaagaatcataatggggaagg
ccatccagcctcgcgtcggggagctttttgcaaaagcctaggcctccaaaaaagcctcctcactacttctggaatagctcagagg
ccgaggcggcctcggcctctgcataaataaaaaaaattagtcagccatg

Vector name: Tat1b

agcttggcccattgcatacgttgtatccatatcataatatgtacatttatattggctcatgtccaacattaccgccatgttgacattgatt
attgactagttattaatagtaatcaattacggggtcattagttcatagcccatatatggagttccgcgttacataacttacggtaaatgg
cccgcctggctgaccgcccaacgacccccgcccattgacgtcaataatgacgtatgttcccatagtaacgccaatagggactttc
cattgacgtcaatgggtggagtatttacggtaaactgcccacttggcagtacatcaagtgtatcatatgccaagtacgccccctatt
gacgtcaatgacggtaaatggcccgcctggcattatgcccagtacatgaccttatgggactttcctacttggcagtacatctacgta
ttagtcatcgctattaccatggtgatgcggttttggcagtacatcaatgggcgtggatagcggtttgactcacggggatttccaagtct
ccaccccattgacgtcaatgggagtttgttttggcaccaaaatcaacgggactttccaaaatgtcgtaacaactccgccccattga
cgcaaatgggcggtaggcgtgtacggtgggaggtctatataagcagagctcgtttagtgaaccgtcagatcgcctggagacgcc
atccacgctgttttgacctccatagaagacaccgggaccgatccagcctcccctcgaagctgatcctgagaacttcagggtgagt
ctatgggacccttgatgttttctttccccttcttttctatggttaagttcatgtcataggaaggggagaagtaacagggtacacatattga
ccaaatcagggtaattttgcatttgtaattttaaaaaatgctttcttcttttaatatactttttttgtttatcttatttctaatactttccctaatctct
ttctttcagggcaataatgatacaatgtatcatgcctctttgcaccattctaaagaataacagtgataatttctgggttaaggcaatagc
aatatttctgcatataaatatttctgcatataaattgtaactgatgtaagaggtttcatattgctaatagcagctacaatccagctaccatt
ctgcttttattttatggttgggataaggctggattattctgagtccaagctaggcccttttgctaatcatgttcatacctcttatcttcctcc
cacagctcctgggcaacgtgctggtctgtgtgctggcccatcactttggcaaagaattccgcgggcggccgcgaaatggagcc
agtagatcctagactagagccctggaagcatccaggaagtcagcctaaaactgcttgtaccacttgctattgtaaaaagtgttgctt
tcattgccaagtttgtttcacaacaaaagccttaggcatctcctatggcaggaagaagcggagacagcgacgaagacctcctca
aggcagtcagactcatcaagtttctctatcaaagcaacccacctcccaaccccgaggggacccgacaggcccgaaggaatag
gatccaagcttatcgataccgtcgacctcgagggcccagatctaattcaccccaccagtgcaggctgcctatcagaaagtggtg
gctggtgtggctaatgccctggcccacaagtatcactaagctcgctttcttgctgtccaatttctattaaaggttcctttgttccctaagt
ccaactactaaactgggggatattatgaagggccttgagcatctggattctgcctaataaaaaacatttattttcattgcaatgatgtat
ttaaattatttctgaatattttactaaaaagggaatgtgggaggtcagtgcatttaaaacataaagaaatgaagagctagttcaaacct
tgggaaaatacactatatcttaaactccatgaaagaaggtgaggctgcaaacagctaatgcacattggcaacagcccctgatgcc
tatgccttattcatccctcagaaaaggattcaagtagaggcttgatttggaggttaaagttttgctatgctgtattttacattacttattgttt
tagctgtcctcatgaatgtcttttcactacccatttgcttatcctgcatctctcagccttgactccactcagttctcttgcttagagatacc
acctttcccctgaagtgttccttccatgttttacggcgagatggtttctcctcgcctggccactcagccttagttgtctctgttgtcttata
gaggtctacttgaagaaggaaaaacagggggcatggtttgactgtcctgtgagcccttcttccctgcctcccccactcacagtga
cccggaatccctcgacatggcagtctagatcattcttgaagacgaaagggcctcgtgatacgcctatttttataggttaatgtcatga
taataatggtttcttagacgtcaggtggcacttttcggggaaatgtgcgcggaacccctatttgtttatttttctaaatacattcaaatat
gtatccgctcatgagacaataaccctgataaatgcttcaataatattgaaaaaggaagagtatgagtattcaacatttccgtgtcgcc
cttattcccttttttgcggcattttgccttcctgtttttgctcacccagaaacgctggtgaaagtaaaagatgctgaagatcagttgggtg

cacgagtgggttacatcgaactggatctcaacagcggtaagatccttgagagttttcgccccgaagaacgttttccaatgatgagc
acttttaaagttctgctatgtggcgcggtattatcccgtattgacgccgggcaagagcaactcggtcgccgcatacactattctcag
aatgacttggttgagtactcaccagtcacagaaaagcatcttacggatggcatgacagtaagagaattatgcagtgctgccataac
catgagtgataacactgcggccaacttacttctgacaacgatcggaggaccgaaggagctaaccgcttttttgcacaacatgggg
gatcatgtaactcgccttgatcgttgggaaccggagctgaatgaagccataccaaacgacgagcgtgacaccacgatgcctgta
gcaatggcaacaacgttgcgcaaactattaactggcgaactacttactctagcttcccggcaacaattaatagactggatggagg
cggataaagttgcaggaccacttctgcgctcggcccttccggctggctggtttattgctgataaatctggagccggtgagcgtggg
tctcgcggtatcattgcagcactggggccagatggtaagccctcccgtatcgtagttatctacacgacggggagtcaggcaacta
tggatgaacgaaatagacagatcgctgagataggtgcctcactgattaagcattggtaactgtcagaccaagtttactcatatatac
tttagattgatttaaaacttcatttttaatttaaaaggatctaggtgaagatcctttttgataatctcatgaccaaaatcccttaacgtgagt
tttcgttccactgagcgtcagaccccgtagaaaagatcaaaggatcttcttgagatcctttttttctgcgcgtaatctgctgcttgcaa
acaaaaaaaccaccgctaccagcggtggtttgtttgccggatcaagagctaccaactctttttccgaaggtaactggcttcagcag
agcgcagataccaaatactgttcttctagtgtagccgtagttaggccaccacttcaagaactctgtagcaccgcctacatacctcg
ctctgctaatcctgttaccagtggctgctgccagtggcgataagtcgtgtcttaccgggttggactcaagacgatagttaccggata
aggcgcagcggtcgggctgaacggggggttcgtgcacacagcccagcttggagcgaacgacctacaccgaactgagatacc
tacagcgtgagctatgagaaagcgccacgcttcccgaagggagaaaggcggacaggtatccggtaagcggcagggtcggaa
caggagagcgcacgagggagcttccagggggaaacgcctggtatctttatagtcctgtcgggtttcgccacctctgacttgagcg
tcgatttttgtgatgctcgtcaggggggcggagcctatggaaaaacgccagcaacggagatgcgccgcgtgcggctgctggag
atggcggacgcgatggatatgttctgccaagggttggtttgcgcattcacagttctccgcaagaattgattggctccaattcttggag
tggtgaatccgttagcgaggtgccgccggcttccattcaggtcgaggtggcccggctccatgcaccgcgacgcaacgcgggg
aggcagacaaggtatagggcggcgcctacaatccatgccaacccgttccatgtgctcgccgaggcggcataaatcgccgtga
cgatcagcggtccaatgatcgaagttaggctggtaagagccgcgagcgatccttgaagctgtccctgatggtcgtcatctacctg
cctggacagcatggcctgcaacgcgggcatcccgatgccgccggaagcgagaagaatcataatggggaaggccatccagcc
tcgcgtcggggagctttttgcaaaagcctaggcctccaaaaaagcctcctcactacttctggaatagctcagaggccgaggcgg
cctcggcctctgcataaataaaaaaaattagtcagccatg

Vector name: Rev 1b

Gacggatcgggagatctcccgatcccctatggtcgactctcagtacaatctgctctgatgccgcatagttaagccagtatctgctc
cctgcttgtgtgttggaggtcgctgagtagtgcgcgagcaaaatttaagctacaacaaggcaaggcttgaccgacaattgcatga
agaatctgcttagggttaggcgttttgcgctgcttcgcgatgtacgggccagatatacgcgttgacattgattattgactagttattaat
agtaatcaattacggggtcattagttcatagcccatatatggagttccgcgttacataacttacggtaaatggcccgcctggctgac
cgcccaacgacccccgcccattgacgtcaataatgacgtatgttcccatagtaacgccaatagggactttccattgacgtcaatg
ggtggactatttacggtaaactgcccacttggcagtacatcaagtgtatcatatgccaagtacgccccctattgacgtcaatgacg
gtaaatggcccgcctggcattatgcccagtacatgaccttatgggactttcctacttggcagtacatctacgtattagtcatcgctatt
accatggtgatgcggttttggcagtacatcaatgggcgtggatagcggtttgactcacggggatttccaagtctccaccccattgac
gtcaatgggagtttgttttggcaccaaaatcaacgggactttccaaaatgtcgtaacaactccgccccattgacgcaaatgggcgg
taggcgtgtacggtgggaggtctatataagcagagctctctggctaactagagaacccactgcttaactggcttatcgaaattaata
cgactcactatagggagacccaagcttggtaccgagctcggatccactagtaacggccgccagtgtgctggaattctgcagatat
ccatcacactggcggccgctcgagcatgcatctagactgccatggcaggaagaagcggagacagcgacgaagacctcctca
aggcagtcagactcatcaagtttctctatcaaagcaacccacctcccaaccccgaggggacccgacaggcccgaaggaatag
aagaagaaggtggagagagagacagagacagatccattcgattagtgaacggatccttagcacttatctgggacgatctgcgga
gcctgtgcctcttcagctaccaccgcttgagagacttactcttgattgtaacgaggattgtggaacttctgggacgcagggggtgg
gaagccctcaaatattggtggaatctcctacagtattggagtcaggaactaaagaataggatccggctattctatagtgtcacctaa
atgctagagctcgctgatcagcctcgactgtgccttctagttgccagccatctgttgtttgcccctcccccgtgccttccttgaccctg
gaaggtgccactcccactgtcctttcctaataaaatgaggaaattgcatcgcattgtctgagtaggtgtcattctattctggggggtg
gggtggggcaggacagcaagggggaggattgggaagacaatagcaggcatgctggggatgcggtgggctctatggcttctga
ggcggaaagaaccagctggggctcgagggggggatccccacgcgccctgtagcggcgcattaagcgcggcgggtgtggtggt
tacgcgcagcgtgaccgctacacttgccagcgccctagcgcccgctcctttcgctttcttcccttcctttctcgccacgttcgccgg

ctttccccgtcaagctctaaatcggggcatccctttagggttccgatttagtgctttacggcacctcgaccccaaaaaacttgattag
ggtgatggttcacgtagtgggccatcgccctgatagacggtttttcgccctttgacgttggagtccacgttctttaatagtggactctt
gttccaaactggaacaacactcaaccctatctcggtctattcttttgatttataagggattttggggatttcggcctattggttaaaaaat
gagctgatttaacaaaaatttaacgcgaattttaacaaaatattaacgtttacaatttaaatatttgcttatacaatcttcctgtttttgggg
cttttctgattatcaaccggggtgggtaccgagctcgaattctgtggaatgtgtgtcagttagggtgtggaaagtccccaggctccc
caggcaggcagaagtatgcaaagcatgcatctcaattagtcagcaaccaggtgtggaaagtccccaggctccccagcaggca
gaagtatgcaaagcatgcatctcaattagtcagcaaccatagtcccgcccctaactccgcccatcccgcccctaactccgccca
gttccgcccattctccgccccatggctgactaattttttttatttatgcagaggccgaggccgcctcggcctctgagctattccagaa
gtagtgaggaggcttttttggaggcctaggcttttgcaaaaagctcccgggagcttggatatccattttcggatctgatcaagagac
aggatgaggatcgtttcgcatgattgaacaagatggattgcacgcaggttctccggccgcttgggtggagaggctattcggctatg
actgggcacaacagacaatcggctgctctgatgccgccgtgttccggctgtcagcgcaggggcgcccggttctttttgtcaagac
cgacctgtccggtgccctgaatgaactgcaggacgaggcagcgcggctatcgtggctggccacgacgggcgttccttgcgca
gctgtgctcgacgttgtcactgaagcgggaagggactggctgctattgggcgaagtgccggggcaggatctcctgtcatctcacc
ttgctcctgccgagaaagtatccatcatggctgatgcaatgcggcggctgcatacgcttgatccggctacctgcccattcgaccac
caagcgaaacatcgcatcgagcgagcacgtactcggatggaagccggtcttgtcgatcaggatgatctggacgaagagcatca
ggggctcgcgccagccgaactgttcgccaggctcaaggcgcgcatgcccgacggcgaggatctcgtcgtgacccatggcgat
gcctgcttgccgaatatcatggtggaaaatggccgcttttctggattcatcgactgtggccggctgggtgtggcggaccgctatca
ggacatagcgttggctacccgtgatattgctgaagagcttggcggcgaatgggctgaccgcttcctcgtgctttacggtatcgccg
ctcccgattcgcagcgcatcgccttctatcgccttcttgacgagttcttctgagcgggactctggggttcgaaatgaccgaccaag
cgacgcccaacctgccatcacgagatttcgattccaccgccgccttctatgaaaggttgggcttcggaatcgttttccgggacgcc
ggctggatgatcctccagcgcggggatctcatgctggagttcttcgcccaccccaacttgtttattgcagcttataatggttacaaat
aaagcaatagcatcacaaatttcacaaataaagcatttttttcactgcattctagttgtggtttgtccaaactcatcaatgtatcttatcat
gtctggatcccgtcgacctcgagagcttggcgtaatcatggtcatagctgtttcctgtgtgaaattgttatccgctcacaattccacac
aacatacgagccggaagcataaagtgtaaagcctggggtgcctaatgagtgagctaactcacattaattgcgttgcgctcactgc
ccgctttccagtcgggaaacctgtcgtgccagctgcattaatgaatcggccaacgcgcggggagaggcggtttgcgtattgggc
gctcttccgcttcctcgctcactgactcgctgcgctcggtcgttcggctgcggcgagcggtatcagctcactcaaaggcggtaata
cggttatccacagaatcaggggataacgcaggaaagaacatgtgagcaaaaggccagcaaaaggccaggaaccgtaaaaa
ggccgcgttgctggcgtttttccataggctccgcccccctgacgagcatcacaaaaatcgacgctcaagtcagaggtggcgaaa
cccgacaggactataaagataccaggcgtttccccctggaagctccctcgtgcgctctcctgttccgaccctgccgcttaccgga
tacctgtccgcctttctcccttcgggaagcgtggcgctttctcaatgctcacgctgtaggtatctcagttcggtgtaggtcgttcgctc
caagctgggctgtgtgcacgaaccccccgttcagcccgaccgctgcgccttatccggtaactatcgtcttgagtccaacccggta
agacacgacttatcgccactggcagcagccactggtaacaggattagcagagcgaggtatgtaggcggtgctacagagttcttg
aagtggtggcctaactacggctacactagaaggacagtatttggtatctgcgctctgctgaagccagttaccttcggaaaaagagt
tggtagctcttgatccggcaaacaaaccaccgctggtagcggtggtttttttgtttgcaagcagcagattacgcgcagaaaaaaag
gatctcaagaagatcctttgatcttttctacggggtctgacgctcagtggaacgaaaactcacgttaagggattttggtcatgagatt
atcaaaaaggatcttcacctagatccttttaaattaaaaatgaagttttaaatcaatctaaagtatatatgagtaaacttggtctgacag
ttaccaatgcttaatcagtgaggcacctatctcagcgatctgtctatttcgttcatccatagttgcctgactccccgtcgtgtagataac
tacgatacgggagggcttaccatctggccccagtgctgcaatgataccgcgagacccacgctcaccggctccagatttatcagc
aataaaccagccagccggaagggccgagcgcagaagtggtcctgcaactttatccgcctccatccagtctattaattgttgccgg
gaagctagagtaagtagttcgccagttaatagtttgcgcaacgttgttgccattgctacaggcatcgtggtgtcacgctcgtcgtttg
gtatggcttcattcagctccggttcccaacgatcaaggcgagttacatgatcccccatgttgtgcaaaaaagcggttagctccttcg
gtcctccgatcgttgtcagaagtaagttggccgcagtgttatcactcatggttatggcagcactgcataattctcttactgtcatgcca
tccgtaagatgcttttctgtgactggtgagtactcaaccaagtcattctgagaatagtgtatgcggcgaccgagttgctcttgcccgg
cgtcaatacgggataataccgcgccacatagcagaactttaaaagtgctcatcattggaaaacgttcttcggggcgaaaactctc
aaggatcttaccgctgttgagatccagttcgatgtaacccactcgtgcacccaactgatcttcagcatcttttactttcaccagcgttt
ctgggtgagcaaaaacaggaaggcaaaatgccgcaaaaaagggaataagggcgacacggaaatgttgaatactcatactcttc
ctttttcaatattattgaagcatttatcagggttattgtctcatgagcggatacatatttgaatgtatttagaaaaataaacaaataggggt
tccgcgcacatttccccgaaaagtgccacctgacgtc

Vector name: Luc2-ZsGreen, variant sequence include CMV promoter, Luciferase and ZsGreen.

tggaagggctaattcactcccaaagaagacaagatatccttgatctgtggatctaccacacacaaggctacttccctgattagcag
aactacacaccagggccaggggtcagatatccactgacctttggatggtgctacaagctagtaccagttgagccagataaggta
gaagaggccaataaaggagagaacaccagcttgttacaccctgtgagcctgcatgggatggatgacccggagagagaagtgtt
agagtggaggtttgacagccgcctagcatttcatcacgtggcccgagagctgcatccggagtacttcaagaactgctgatatcga
gcttgctacaagggactttccgctggggactttccagggaggcgtggcctgggcgggactggggagtggcgagccctcagatc
ctgcatataagcagctgctttttgcctgtactgggtctctctggttagaccagatctgagcctgggagctctctggctaactagggaa
cccactgcttaagcctcaataaagcttgccttgagtgcttcaagtagtgtgtgcccgtctgttgtgtgactctggtaactagagatcc
ctcagacccttttagtcagtgtggaaaatctctagcagtggcgcccgaacagggacttgaaagcgaaagggaaaccagaggag
ctctctcgacgcaggactcggcttgctgaagcgcgcacggcaagaggcgaggggcggcgactggtgagtacgccaaaaatttt
gactagcggaggctagaaggagagagatgggtgcgagagcgtcagtattaagcgggggagaattagatcgcgatgggaaaa
aattcggttaaggccagggggaaagaaaaaatataaattaaaacatatagtatgggcaagcagggagctagaacgattcgcagt
taatcctggcctgttagaaacatcagaaggctgtagacaaatactgggacagctacaaccatcccttcagacaggatcagaaga
acttagatcattatataatacagtagcaaccctctattgtgtgcatcaaaggatagagataaaagacaccaaggaagctttagaca
agatagaggaagagcaaaacaaaagtaagaccaccgcacagcaagcggccggccgctgatcttcagacctggaggaggag
atatgagggacaattggagaagtgaattatataaatataaagtagtaaaaattgaaccattaggagtagcacccaccaaggcaaa
gagaagagtggtgcagagagaaaaaagagcagtgggaataggagctttgttccttgggttcttgggagcagcaggaagcactat
gggcgcagcgtcaatgacgctgacggtacaggccagacaattattgtctggtatagtgcagcagcagaacaatttgctgagggc
tattgaggcgcaacagcatctgttgcaactcacagtctggggcatcaagcagctccaggcaagaatcctggctgtggaaagata
cctaaaggatcaacagctcctggggatttggggttgctctggaaaactcatttgcaccactgctgtgccttggaatgctagttggag
taataaatctctggaacagatttggaatcacacgacctggatggagtgggacagagaaattaacaattacacaagcttaatacact
ccttaattgaagaatcgcaaaaccagcaagaaaagaatgaacaagaattattggaattagataaatgggcaagtttgtggaattgg
tttaacataacaaattggctgtggtatataaaattattcataatgatagtaggaggcttggtaggtttaagaatagtttttgctgtactttct
atagtgaatagagttaggcagggatattcaccattatcgtttcagacccacctcccaaccccgaggggacccgacaggcccgaa
ggaatagaagaagaaggtggagagagagacagagacagatccattcgattagtgaacggatctcgacggtatcgccgaattca
caaatggcagtattcatccacaattttaaaagaaaaggggggattggggggtacagtgcaggggaaagaatagtagacataata
gcaacagacatacaaactaaagaattacaaaaacaaattacaaaaattcaaaattttcgggtttattacagggacagcagagatc
cagtttggactagtggagttccgcgttacataacttacggtaaatggcccgcctggctgaccgcccaacgacccccgcccattga
cgtcaataatgacgtatgttcccatagtaacgccaatagggactttccattgacgtcaatgggtggagtatttacggtaaactgccc
acttggcagtacatcaagtgtatcatatgccaagtacgccccctattgacgtcaatgacggtaaatggcccgcctggcattatgcc
cagtacatgaccttatgggactttcctacttggcagtacatctacgtattagtcatcgctattaccatggtgatgcggttttggcagtac
atcaatgggcgtggatagcggtttgactcacggggatttccaagtctccaccccattgacgtcaatgggagtttgttttggcaccaa
aatcaacgggactttccaaaatgtcgtaacaactccgccccattgacgcaaatgggcggtaggcgtgtacggtgggaggtctata
taagcagagctcgtttagtgaaccgtcagatcgcctggagacgccatccacgctgttttgacctccatagaagacaccggcggc
cgccatggaagatgccaaaaacattaagaagggcccagcgccattctacccactcgaagacgggaccgccggcgagcagct
gcacaaagccatgaagcgctacgccctggtgcccggcaccatcgcctttaccgacgcacatatcgaggtggacattacctacg
ccgagtacttcgagatgagcgttcggctggcagaagctatgaagcgctatgggctgaatacaaaccatcggatcgtggtgtgca
gcgagaatagcttgcagttcttcatgcccgtgttgggtgccctgttcatcggtgtggctgtggccccagctaacgacatctacaacg
agcgcgagctgctgaacagcatgggcatcagccagcccaccgtcgtattcgtgagcaagaaagggctgcaaaagatcctcaa
cgtgcaaaagaagctaccgatcatacaaaagatcatcatcatggatagcaagaccgactaccagggcttccaaagcatgtaca
ccttcgtgacttcccatttgccacccggcttcaacgagtacgacttcgtgcccgagagcttcgaccgggacaaaaccatcgccct
gatcatgaacagtagtggcagtaccggattgcccaagggcgtagccctaccgcaccgcaccgcttgtgtccgattcagtcatgc
ccgcgaccccatcttcggcaaccagatcatccccgacaccgctatcctcagcgtggtgccatttcaccacggcttcggcatgttc
accacgctgggctacttgatctgcggctttcgggtcgtgctcatgtaccgcttcgaggaggagctattcttgcgcagcttgcaaga
ctataagattcaatctgccctgctggtgcccacactatttagcttcttcgctaagagcactctcatcgacaagtacgacctaagcaa
cttgcacgagatcgccagcggcggggcgccgctcagcaaggaggtaggtgaggccgtggccaaacgcttccacctaccagg
catccgccagggctacggcctgacagaaacaaccagcgccattctgatcacccccgaaggggacgacaagcctggcgcagt

aggcaaggtggtgcccttcttcgaggctaaggtggtggacttggacaccggtaagacactgggtgtgaaccagcgcggcgagc
tgtgcgtccgtggccccatgatcatgagcggctacgttaacaaccccgaggctacaaacgctctcatcgacaaggacggctgg
ctgcacagcggcgacatcgcctactgggacgaggacgagcacttcttcatcgtggaccggctgaagagcctgatcaaatacaa
gggctaccaggtagccccagccgaactggagagcatcctgctgcaacaccccaacatcttcgacgccggggtcgccggcctg
cccgacgacgatgccggcgagctgcccgccgcagtcgtcgtgctggaacacggtaaaaccatgaccgagaaggagatcgtg
gactatgtggccagccaggttacaaccgccaagaagctgcgcggtggtgttgtgttcgtggacgaggtgcctaaaggactgacc
ggcaagttggacgcccgcaagatccgcgagattctcattaaggccaagaagggcggcaagatcgccgtgtaaaggatccctc
cccccccccctaacgttactggccgaagccgcttggaataaggccggtgtgcgtttgtctatatgttattttccaccatattgccgtctt
ttggcaatgtgagggcccggaaacctggccctgtcttcttgacgagcattcctaggggtctttcccctctcgccaaaggaatgcaa
ggtctgttgaatgtcgtgaaggaagcagttcctctggaagcttcttgaagacaaacaacgtctgtagcgaccctttgcaggcagcg
gaaccccccacctggcgacaggtgcctctgcggccaaaagccacgtgtataagatacacctgcaaaggcggcacaacccca
gtgccacgttgtgagttggatagttgtggaaagagtcaaatggctctcctcaagcgtattcaacaaggggctgaaggatgcccag
aaggtaccccattgtatgggatctgatctggggcctcggtgcacatgctttacatgtgtttagtcgaggttaaaaaaacgtctaggc
cccccgaaccacggggacgtggttttcctttgaaaaacacgatgataatatggccacacatatggcccagtccaagcacggcct
gaccaaggagatgaccatgaagtaccgcatggagggctgcgtggacggccacaagttcgtgatcaccggcgagggcatcgg
ctaccccttcaagggcaagcaggccatcaacctgtgcgtggtggagggcggccccttgcccttcgccgaggacatcttgtccgc
cgccttcatgtacggcaaccgcgtgttcaccgagtaccccaggacatcgtcgactacttcaagaactcctgccccgccggcta
cacctgggaccgctccttcctgttcgaggacggcgccgtgtgcatctgcaacgccgacatcaccgtgagcgtggaggagaact
gcatgtaccacgagtccaagttctacggcgtgaacttccccgccgacggccccgtgatgaagaagatgaccgacaactggga
gccctcctgcgagaagatcatccccgtgcccaagcagggcatcttgaagggcgacgtgagcatgtacctgctgctgaaggacg
gtggccgcttgcgctgccagttcgacaccgtgtacaaggccaagtccgtgccccgcaagatgcccgactggcacttcatccag
cacaagctgacccgcgaggaccgcagcgacgccaagaaccagaagtggcacctgaccgagcacgccatcgcctccggctc
cgccttgccctgaatcgatagatcctaatcaacctctggattacaaaatttgtgaaagattgactggtattcttaactatgttgctcctttt
tacgctatgtggatacgctgctttaatgcctttgtatcatgctattgcttcccgtatggctttcattttctcctccttgtataaatcctggttg
ctgtctctttatgaggagttgtggcccgttgtcaggcaacgtggcgtggtgtgcactgtgtttgctgacgcaacccccactggttgg
ggcattgccaccacctgtcagctcctttccgggactttcgctttccccctccctattgccacggcggaactcatcgccgcctgcctt
gcccgctgctggacaggggctcggctgttgggcactgacaattccgtggtgttgtcggggaaatcatcgtcctttccttggctgctc
gcctgtgttgccacctggattctgcgcgggacgtccttctgctacgtcccttcggccctcaatccagcggaccttccttcccgcgg
cctgctgccggctctgcggcctcttccgcgtcttcgccttcgccctcagacgagtcggatctccctttgggccgcctccccgcctg
agatcctttaagaccaatgacttacaaggcagctgtagatcttagccactttttaaaagaaaaggggggactggaagggctaattc
actcccaacgaagacaagatctgctttttgcttgtactgggtctctctggttagaccagatctgagcctgggagctctctggctaact
agggaacccactgcttaagcctcaataaagcttgccttgagtgcttcaagtagtgtgtgcccgtctgttgtgtgactctggtaactag
agatccctcagacccttttagtcagtgtggaaaatctctagcagtagtagttcatgtcatcttattattcagtatttataacttgcaaaga
aatgaatatcagagagtgagaggcccgggttaattaaggaaagggctagatcattcttgaagacgaaagggcctcgtgatacgc
ctatttttataggttaatgtcatgataataatggtttcttagacgtcaggtggcacttttcggggaaatgtgcgcggaacccctatttgttt
atttttctaaatacattcaaatatgtatccgctcatgagacaataaccctgataaatgcttcaataatattgaaaaaggaagagtatga
gtattcaacatttccgtgtcgcccttattcccttttttgcggcattttgccttcctgtttttgctcacccagaaacgctggtgaaagtaaaa
gatgctgaagatcagttgggtgcacgagtgggttacatcgaactggatctcaacagcggtaagatccttgagagttttcgccccga
agaacgttttccaatgatgagcacttttaaagttctgctatgtggcgcggtattatcccgtgttgacgccgggcaagagcaactcgg
tcgccgcatacactattctcagaatgacttggttgagtactcaccagtcacagaaaagcatcttacggatggcatgacagtaagag
aattatgcagtgctgccataaccatgagtgataacactgcggccaacttacttctgacaacgatcggaggaccgaaggagctaa
ccgcttttttgcacaacatgggggatcatgtaactcgccttgatcgttgggaaccggagctgaatgaagccataccaaacgacga
gcgtgacaccacgatgcctgtagcaatggcaacaacgttgcgcaaactattaactggcgaactacttactctagcttcccggcaa
caattaatagactggatggaggcggataaagttgcaggaccacttctgcgctcggcccttccggctggctggtttattgctgataaa
tctggagccggtgagcgtgggtctcgcggtatcattgcagcactggggccagatggtaagccctcccgtatcgtagttatctacac
gacggggagtcaggcaactatggatgaacgaaatagacagatcgctgagataggtgcctcactgattaagcattggtaactgtc
agaccaagtttactcatatatactttagattgatttaaaacttcatttttaatttaaaaggatctaggtgaagatcctttttgataatctcatg
accaaaatcccttaacgtgagttttcgttccactgagcgtcagaccccgtagaaaagatcaaaggatcttcttgagatcctttttttct
gcgcgtaatctgctgcttgcaaacaaaaaaaccaccgctaccagcggtggtttgtttgccggatcaagagctaccaactctttttcc

gaaggtaactggcttcagcagagcgcagataccaaatactgttcttctagtgtagccgtagttaggccaccacttcaagaactctg
tagcaccgcctacatacctcgctctgctaatcctgttaccagtggctgctgccagtggcgataagtcgtgtcttaccgggttggact
caagacgatagttaccggataaggcgcagcggtcgggctgaacggggggttcgtgcacacagcccagcttggagcgaacga
cctacaccgaactgagatacctacagcgtgagctatgagaaagcgccacgcttcccgaagggagaaaggcggacaggtatcc
ggtaagcggcagggtcggaacaggagagcgcacgagggagcttccagggggaaacgcctggtatctttatagtcctgtcgggt
ttcgccacctctgacttgagcgtcgatttttgtgatgctcgtcaggggggcggagcctatggaaaaacgccagcaacgcggccttt
ttacggttcctggccttttgctggccttttgctcacatgttctttcctgcgttatcccctgattctgtggataaccgtattaccgcctttgag
tgagctgataccgctcgccgcagccgaacgaccgagcgcagcgagtcagtgagcgaggaagcggaagagcgcccaatacg
caaaccgcctctccccgcgcgttggccgattcattaatgcagcaagctcatggctgactaatttttttttatttatgcagaggccgagg
ccgcctcggcctctgagctattccagaagtagtgaggaggcttttttggaggcctaggcttttgcaaaaagctccccgtggcacga
caggtttcccgactggaaagcgggcagtgagcgcaacgcaattaatgtgagttagctcactcattaggcaccccaggctttacac
tttatgcttccggctcgtatgttgtgtggaattgtgagcggataacaatttcacacaggaaacagctatgacatgattacgaatttcac
aaataaagcatttttttcactgcattctagttgtggtttgtccaaactcatcaatgtatcttatcatgtctggatcaactggataactcaag
ctaaccaaaatcatcccaaacttcccaccccataccctattaccactgccaattacctgtggtttcatttactctaaacctgtgattcc
tctgaattattttcattttaaagaaattgtatttgttaaatatgtactacaaacttagtagt.

Plasmid to increase spike protein infectivity was prepared by Gibson assembly. Vector name: Spike G614 Δ19. The capital sequence was the modified sequence from wide type Spike.

agcttggcccattgcatacgttgtatccatatcataatatgtacatttatattggctcatgtccaacattaccgccatgttgacattgatt
attgactagttattaatagtaatcaattacggggtcattagttcatagcccatatatggagttccgcgttacataacttacggtaaatgg
cccgcctggctgaccgcccaacgacccccgcccattgacgtcaataatgacgtatgttcccatagtaacgccaatagggactttc
cattgacgtcaatgggtggagtatttacggtaaactgcccacttggcagtacatcaagtgtatcatatgccaagtacgccccctatt
gacgtcaatgacggtaaatggcccgcctggcattatgcccagtacatgaccttatgggactttcctacttggcagtacatctacgta
ttagtcatcgctattaccatggtgatgcggttttggcagtacatcaatgggcgtggatagcggtttgactcacggggatttccaagtct
ccaccccattgacgtcaatgggagtttgttttggcaccaaaatcaacgggactttccaaaatgtcgtaacaactccgccccattga
cgcaaatgggcggtaggcgtgtacggtgggaggtctatataagcagagctcgtttagtgaaccgtcagatcgcctggagacgcc
atccacgctgttttgacctccatagaagacaccgggaccgatccagcctcccctcgaagctgatcctgagaacttcagggtgagt
ctatgggacccttgatgttttctttccccttcttttctatggttaagttcatgtcataggaaggggagaagtaacagggtacacatattga
ccaaatcagggtaattttgcatttgtaattttaaaaaatgctttcttcttttaatatacttttttgtttatcttatttctaatactttccctaatctct
ttctttcagggcaataatgatacaatgtatcatgcctctttgcaccattctaaagaataacagtgataatttctgggttaaggcaatagc
aatatttctgcatataaatatttctgcatataaattgtaactgatgtaagaggtttcatattgctaatagcagctacaatccagctaccatt
ctgcttttattttatggttgggataaggctggattattctgagtccaagctaggcccttttgctaatcatgttcatacctcttatcttcctcc
cacagctcctgggcaacgtgctggtctgtgtgctggcccatcactttggcaaagaattccgcgggcggccgccATGTTCGT
GTTCCTGGTACTCCTTCCTTTGGTGTCTTCCCAGTGTGTAAATCTTACCACTCGGACCCAGCT
TCCACCCGCCTACACCAACAGTTTCACACGCGGCGTCTACTATCCTGACAAGGTGTTTAGGA
GTTCAGTCTTGCACTCAACTCAAGACTTGTTCCTCCCTTTCTTTAGCAATGTGACGTGGTTTCA
TGCCATTCATGTCTCCGGCACAAACGGAACGAAGCGCTTTGATAATCCTGTGCTCCCGTTCAA
CGATGGAGTGTACTTCGCGTCCACAGAGAAGAGCAATATCATTCGAGGTTGGATCTTCGGAA
CGACACTCGACTCAAAGACGCAGTCCCTTCTCATCGTCAATAATGCCACGAACGTGGTCATC
AAAGTGTGCGAGTTTCAATTCTGTAATGATCCCTTCCTGGGCGTCTATTATCACAAGAACAACA
AATCCTGGATGGAGTCCGAATTTAGAGTCTACTCCAGCGCCAACAACTGCACTTTCGAATACG
TATCACAGCCATTCTTGATGGACCTTGAAGGAAAGCAGGGTAATTTCAAGAACTTGAGGGAGT
TCGTATTCAAGAATATCGACGGGTACTTTAAGATTTATAGCAAACACACACCCATTAATTTGGT
GCGGGATCTTCCTCAGGGATTTAGTGCTCTTGAGCCTCTCGTTGACCTCCCTATTGGCATTAA
CATCACCCGCTTTCAAACCCTGTTGGCCCTGCATCGGTCCTACCTGACACCGGGCGACTCAA
GTTCCGGATGGACCGCAGGTGCCGCCGCATACTATGTGGGCTACCTTCAGCCAAGAACATTT
CTGCTGAAATATAATGAGAACGGGACCATTACAGATGCGGTGGATTGTGCACTCGACCCTCT
GTCTGAGACGAAATGCACCCTTAAGAGCTTCACGGTGGAGAAAGGCATTTATCAGACTTCTAA

CTTCAGAGTTCAACCCACCGAGTCCATTGTGCGATTCCCAAATATTACGAATTTGTGCCCATTT
GGTGAGGTCTTCAATGCTACTCGATTCGCCTCAGTTTATGCATGGAACCGAAAGAGAATTTCC
AATTGTGTGGCGGACTACTCAGTATTGTATAATAGTGCAAGCTTTAGCACATTCAAATGTTACG
GCGTGTCTCCAACGAAGCTGAACGATCTCTGTTTCACAAACGTTTATGCGGATTCCTTCGTGA
TTCGCGGCGATGAGGTCCGACAGATTGCGCCTGGGCAAACGGGTAAGATCGCTGATTACAA
CTATAAGTTGCCGGACGATTTCACAGGATGTGTCATAGCTTGGAATAGCAATAATTTGGACAG
TAAGGTTGGCGGAAACTACAATTATTTGTACAGGTTGTTTCGCAAGTCAAATTTGAAACCATTT
GAGAGAGATATATCTACGGAGATATATCAAGCCGGCTCTACACCATGTAATGGTGTGGAGGG
CTTTAACTGCTACTTTCCACTCCAGTCATATGGTTTCCAACCTACAAATGGAGTAGGGTATCAA
CCGTACAGAGTTGTGGTCTTGAGTTTCGAATTGCTCCACGCTCCAGCAACGGTATGCGGTCC
TAAGAAATCCACAAATCTTGTGAAGAACAAGTGCGTAAATTTCAACTTCAATGGGCTGACTGGA
ACAGGCGTGCTGACTGAGAGTAACAAGAAGTTCTTGCCTTTCCAACAATTCGGGCGGGACAT
AGCTGATACCACTGACGCCGTCCGCGACCCTCAGACCCTGGAGATTCTGGACATAACTCCTT
GTTCTTTCGGTGGCGTCAGTGTTATCACTCCCGGGACCAACACCTCCAACCAAGTCGCGGTC
CTCTATCAAGGCGTCAACTGTACGGAAGTACCGGTAGCCATCCATGCGGACCAACTTACACC
GACTTGGAGGGTTTACTCTACAGGAAGCAATGTCTTTCAAACACGAGCCGGGTGTCTGATCG
GAGCAGAACACGTTAACAACAGCTACGAATGTGACATACCAATAGGCGCAGGGATTTGTGCT
TCATATCAGACACAGACCAATAGCCCGagcAGAGCGAGtAGCGTAGCAAGCCAAAGCATCATC
GCGTACACGATGAGCCTCGGAGCAGAGAACAGCGTCGCGTATAGCAATAATTCAATAGCTAT
CCCAACAAATTTCACTATTTCGGTCACCACTGAGATTCTGCCGGTCTCCATGACCAAGACATC
CGTCGATTGTACTATGTACATATGCGGCGACAGCACGGAGTGCAGTAACTTGCTCCTTCAGTA
CGGTTCCTTCTGTACGCAGCTTAACCGGGCACTGACGGGTATCGCGGTAGAACAGGACAAG
AACACACAGGAGGTCTTCGCGCAGGTCAAACAAATCTACAAGACACCACCCATAAAGGACTT
CGGCGGGTTCAATTTCAGCCAAATCCTGCCGGACCCTTCCAAACCTAGTAAGAGGTCATTCA
TTGAGGATCTTCTGTTTAACAAAGTTACGCTTGCGGACGCGGGATTCATTAAGCAGTATGGTG
ACTGCCTTGGAGATATTGCCGCCAGGGATTTGATATGTGCACAGAAATTTAACGGCCTCACCG
TTCTGCCGCCTCTGCTCACCGATGAGATGATAGCGCAGTACACGAGCGCACTCCTGGCAGG
TACAATTACAAGCGGATGGACATTCGGTGCAGGAGCAGCGTTGCAGATACCCTTTGCTATGC
AGATGGCTTATCGATTTAACGGGATTGGCGTCACGCAGAACGTCCTTTATGAGAATCAGAAAT
TGATTGCAAATCAGTTCAATAGTGCTATCGGTAAGATTCAGGACAGCTTGAGCAGTACCGCGT
CTGCACTGGGAAAGTTGCAGGACGTGGTGAATCAGAATGCACAAGCACTGAATACCTTGGTT
AAGCAATTGAGTAGCAATTTCGGCGCCATATCAAGTGTACTGAATGATATCCTGTCACGGTTG
GACAAGGTAGAAGCCGAAGTTCAGATTGACCGCTTGATCACCGGGCGCCTCCAAAGTCTGCA
GACCTACGTCACACAACAATTGATCAGAGCAGCAGAGATAAGAGCATCTGCTAACCTGGCTG
CCACTAAGATGTCTGAATGTGTGCTTGGGCAGTCAAAGAGGGTAGATTTCTGCGGAAAGGGC
TACCACCTTATGTCTTTCCCTCAGAGCGCTCCGCATGGTGTGGTCTTTCTCCATGTGACTTAT
GTGCCTGCTCAAGAGAAGAACTTTACGACGGCGCCCGCTATATGCCATGATGGTAAGGCGCA
CTTTCCAAGGGAGGGAGTGTTCGTGTCCAACGGCACTCACTGGTTTGTCACCCAACGAAATTT
CTACGAGCCTCAAATTATTACCACCGACAATACCTTTGTTAGCGGTAACTGTGACGTCGTAATT
GGGATTGTTAATAATACAGTCTACGATCCTCTGCAGCCGGAACTGGACTCCTTTAAAGAGGAG
CTGGACAAATATTTCAAGAACCACACATCTCCTGACGTAGATCTTGGAGACATAAGCGGTATA
AATGCAAGTGTTGTTAACATTCAGAAAGAAATAGATAGGTTGAACGAAGTTGCGAAGAACCTTA
ACGAGTCACTGATAGACCTCCAAGAGCTTGGGAAGTACGAGCAATATATCAAGTGGCCTTGG
TATATTTGGCTCGGGTTCATAGCAGGACTTATCGCTATAGTCATGGTGACTATAATGCTGTGCT
GCATGACAAGCTGCTGCAGCTGTCTCAAAGGCTGTTGCTCTTGCGGCTCTTGCTGCTAATGAa
agcttatcgataccgtcgacctcgagggcccagatctaattcaccccaccagtgcaggctgcctatcagaaagtggtggctggtg
tggctaatgccctggcccacaagtatcactaagctcgctttcttgctgtccaatttctattaaaggttcctttgttccctaagtccaacta
ctaaactggggatattatgaagggccttgagcatctggattctgcctaataaaaaacatttattttcattgcaatgatgtatttaaatta
tttctgaatattttactaaaaagggaatgtgggaggtcagtgcatttaaaacataaagaaatgaagagctagttcaaaccttgggaa

aatacactatatcttaaactccatgaaagaaggtgaggctgcaaacagctaatgcacattggcaacagcccctgatgcctatgcc
ttattcatccctcagaaaaggattcaagtagaggcttgatttggaggttaaagttttgctatgctgtattttacattacttattgttttagctg
tcctcatgaatgtcttttcactacccatttgcttatcctgcatctctcagccttgactccactcagttctcttgcttagagataccacctttc
ccctgaagtgttccttccatgttttacggcgagatggtttctcctcgcctggccactcagccttagttgtctctgttgtcttatagaggtc
tacttgaagaaggaaaaacagggggcatggtttgactgtcctgtgagcccttcttccctgcctcccccactcacagtgacccgga
atccctcgacatggcagtctagatcattcttgaagacgaaagggcctcgtgatacgcctatttttataggttaatgtcatgataataat
ggtttcttagacgtcaggtggcacttttcggggaaatgtgcgcggaacccctatttgtttatttttctaaatacattcaaatatgtatccg
ctcatgagacaataaccctgataaatgcttcaataatattgaaaaaggaagagtatgagtattcaacatttccgtgtcgcccttattcc
cttttttgcggcattttgccttcctgtttttgctcacccagaaacgctggtgaaagtaaaagatgctgaagatcagttgggtgcacgag
tgggttacatcgaactggatctcaacagcggtaagatccttgagagttttcgccccgaagaacgttttccaatgatgagcacttttaa
agttctgctatgtggcgcggtattatcccgtattgacgccgggcaagagcaactcggtcgccgcatacactattctcagaatgactt
ggttgagtactcaccagtcacagaaaagcatcttacggatggcatgacagtaagagaattatgcagtgctgccataaccatgagt
gataacactgcggccaacttacttctgacaacgatcggaggaccgaaggagctaaccgcttttttgcacaacatgggggatcatg
taactcgccttgatcgttgggaaccggagctgaatgaagccataccaaacgacgagcgtgacaccacgatgcctgtagcaatg
gcaacaacgttgcgcaaactattaactggcgaactacttactctagcttcccggcaacaattaatagactggatggaggcggata
aagttgcaggaccacttctgcgctcggcccttccggctggctggtttattgctgataaatctggagccggtgagcgtgggtctcgc
ggtatcattgcagcactggggccagatggtaagccctcccgtatcgtagttatctacacgacggggagtcaggcaactatggatg
aacgaaatagacagatcgctgagataggtgcctcactgattaagcattggtaactgtcagaccaagtttactcatatatactttagat
tgatttaaaacttcatttttaatttaaaaggatctaggtgaagatcctttttgataatctcatgaccaaaatcccttaacgtgagttttcgtt
ccactgagcgtcagaccccgtagaaaagatcaaaggatcttcttgagatcctttttttctgcgcgtaatctgctgcttgcaaacaaa
aaaaccaccgctaccagcggtggtttgtttgccggatcaagagctaccaactctttttccgaaggtaactggcttcagcagagcgc
agataccaaatactgttcttctagtgtagccgtagttaggccaccacttcaagaactctgtagcaccgcctacatacctcgctctgct
aatcctgttaccagtggctgctgccagtggcgataagtcgtgtcttaccgggttggactcaagacgatagttaccggataaggcgc
agcggtcgggctgaacggggggttcgtgcacacagcccagcttggagcgaacgacctacaccgaactgagatacctacagc
gtgagctatgagaaagcgccacgcttcccgaagggagaaaggcggacaggtatccggtaagcggcagggtcggaacagga
gagcgcacgagggagcttccagggggaaacgcctggtatctttatagtcctgtcgggtttcgccacctctgacttgagcgtcgatt
tttgtgatgctcgtcaggggggcggagcctatggaaaaacgccagcaacggatgcgccgcgtgcggctgctggagatggcgg
acgcgatggatatgttctgccaagggttggtttgcgcattcacagttctccgcaagaattgattggctccaattcttggagtggtgaat
ccgttagcgaggtgccgccggcttccattcaggtcgaggtggcccggctccatgcaccgcgacgcaacgcggggaggcaga
caaggtatagggcggcgcctacaatccatgccaacccgttccatgtgctcgccgaggcggcataaatccccgtgacgatcagc
ggtccaatgatcgaagttaggctggtaagagccgcgagcgatccttgaagctgtccctgatggtcgtcatctacctgcctggaca
gcatggcctgcaacgcgggcatcccgatgccgccggaagcgagaagaatcataatggggaaggccatccagcctcgcgtcg
gggagctttttgcaaaagcctaggcctccaaaaaagcctcctcactacttctggaatagctcagaggccgaggcggcctcggcc
tctgcataaataaaaaaaattagtcagccatg.

Plasmids to insert N gene of SARS-COV-2 by Gibson assembly, vector name: CoV2Ngene-Luc2-ZsGreen. Variant sequences include the N gene and ZsGreen sequence. The capital sequence was the inserted N gene sequence.

tggaagggctaattcactcccaaagaagacaagatatccttgatctgtggatctaccacacacaaggctacttccctgattagcag
aactacacaccagggccaggggtcagatatccactgacctttggatggtgctacaagctagtaccagttgagccagataaggta
gaagaggccaataaaggagagaacaccagcttgttacaccctgtgagcctgcatgggatggatgacccggagagagaagtgtt
agagtggaggtttgacagccgcctagcatttcatcacgtggcccgagagctgcatccggagtacttcaagaactgctgatatcga
gcttgctacaagggactttccgctggggactttccagggaggcgtggcctgggcgggactggggagtggcgagccctcagatc
ctgcatataagcagctgctttttgcctgtactgggtctctctggttagaccagatctgagcctgggagctctctggctaactagggaa
cccactgcttaagcctcaataaagcttgccttgagtgcttcaagtagtgtgtgcccgtctgttgtgtgactctggtaactagagatcc
ctcagacccttttagtcagtgtggaaaatctctagcagtggcgcccgaacagggacttgaaagcgaaagggaaaccagaggag
ctctctcgacgcaggactcggcttgctgaagcgcgcacggcaagaggcgaggggcggcgactggtgagtacgccaaaaatttt
gactagcggaggctagaaggagagagatgggtgcgagagcgtcagtattaagcgggggagaattagatcgcgatgggaaaa

aattcggttaaggccagggggaaagaaaaaatataaattaaaacatatagtatgggcaagcagggagctagaacgattcgcagt
taatcctggcctgttagaaacatcagaaggctgtagacaaatactgggacagctacaaccatcccttcagacaggatcagaaga
acttagatcattatataatacagtagcaaccctctattgtgtgcatcaaaggatagagataaaagacaccaaggaagctttagaca
agatagaggaagagcaaaacaaaagtaagaccaccgcacagcaagcggccggccgctgatcttcagacctggaggaggag
atatgagggacaattggagaagtgaattatataaatataaagtagtaaaaattgaaccattaggagtagcacccaccaaggcaaa
gagaagagtggtgcagagagaaaaaagagcagtgggaataggagctttgttccttgggttcttgggagcagcaggaagcactat
gggcgcagcgtcaatgacgctgacggtacaggccagacaattattgtctggtatagtgcagcagcagaacaatttgctgagggc
tattgaggcgcaacagcatctgttgcaactcacagtctggggcatcaagcagctccaggcaagaatcctggctgtggaaagata
cctaaaggatcaacagctcctggggatttggggttgctctggaaaactcatttgcaccactgctgtgccttggaatgctagttggag
taataaatctctggaacagatttggaatcacacgacctggatggagtgggacagagaaattaacaattacacaagcttaatacact
ccttaattgaagaatcgcaaaaccagcaagaaaagaatgaacaagaattattggaattagataaatgggcaagtttgtggaattgg
tttaacataacaaattggctgtggtatataaaattattcataatgatagtaggaggcttggtaggtttaagaatagtttttgctgtactttct
atagtgaatagagttaggcagggatattcaccattatcgtttcagacccacctcccaaccccgaggggacccgacaggcccgaa
ggaatagaagaagaaggtggagagagagacagagacagatccattcgattagtgaacggatctcgacggtatcgccgaattca
caaatggcagtattcatccacaattttaaaagaaaaggggggattggggggtacagtgcaggggaaagaatagtagacataata
gcaacagacatacaaactaaagaattacaaaaacaaattacaaaaattcaaaattttcgggtttattacagggacagcagagatc
cagtttggactagtggagttccgcgttacataacttacggtaaatggcccgcctggctgaccgcccaacgacccccgcccattga
cgtcaataatgacgtatgttcccatagtaacgccaatagggactttccattgacgtcaatgggtggagtatttacggtaaactgccc
acttggcagtacatcaagtgtatcatatgccaagtacgccccctattgacgtcaatgacggtaaatggcccgcctggcattatgcc
cagtacatgaccttatgggactttcctacttggcagtacatctacgtattagtcatcgctattaccatggtgatgcggttttggcagtac
atcaatgggcgtggatagcggtttgactcacggggatttccaagtctccaccccattgacgtcaatgggagtttgttttggcaccaa
aatcaacgggactttccaaaatgtcgtaacaactccgccccattgacgcaaatgggcggtaggcgtgtacggtgggaggtctata
taagcagagctcgtttagtgaaccgtcagatcgcctggagacgccatccacgctgttttgacctccatagaagacaccggcggc
cgccATGTCTGATAATGGACCCCAAAATCAGCGAAATGCACCCCGCATTACGTTTGGTGGACC
CTCAGATTCAACTGGCAGTAACCAGAATGGAGAACGCAGTGGGGCGCGATCAAAACAACGTC
GGCCCCAAGGTTTACCCAATAATACTGCGTCTTGGTTCACCGCTCTCACTCAACATGGCAAG
GAAGACCTTAAATTCCCTCGAGGACAAGGCGTTCCAATTAACACCAATAGCAGTCCAGATGAC
CAAATTGGCTACTACCGAAGAGCTACCAGACGAATTCGTGGTGGTGACGGTAAAATGAAAGA
TCTCAGTCCAAGATGGTATTTCTACTACCTAGGAACTGGGCCAGAAGCTGGACTTCCCTATGG
TGCTAACAAAGACGGCATCATATGGGTTGCAACTGAGGGAGCCTTGAATACACCAAAAGATC
ACATTGGCACCCGCAATCCTGCTAACAATGCTGCAATCGTGCTACAACTTCCTCAAGGAACAA
CATTGCCAAAAGGCTTCTACGCAGAAGGGAGCAGAGGCGGCAGTCAAGCCTCTTCTCGTTCC
TCATCACGTAGTCGCAACAGTTCAAGAAATTCAACTCCAGGCAGCAGTAGGGGAACTTCTCCT
GCTAGAATGGCTGGCAATGGCGGTGATGCTGCTCTTGCTTTGCTGCTGCTTGACAGATTGAA
CCAGCTTGAGAGCAAAATGTCTGGTAAAGGCCAACAACAACAAGGCCAAACTGTCACTAAGA
AATCTGCTGCTGAGGCTTCTAAGAAGCCTCGGCAAAAACGTACTGCCACTAAAGCATACAATG
TAACACAAGCTTTCGGCAGACGTGGTCCAGAACAAACCCAAGGAAATTTTGGGGACCAGGAA
CTAATCAGACAAGGAACTGATTACAAACATTGGCCGCAAATTGCACAATTTGCCCCCAGCGCT
TCAGCGTTCTTCGGAATGTCGCGCATTGGCATGGAAGTCACACCTTCGGGAACGTGGTTGAC
CTACACAGGTGCCATCAAATTGGATGACAAAGATCCAAATTTCAAAGATCAAGTCATTTTGCTG
AATAAGCATATTGACGCATACAAAACATTCCCACCAACAGAGCCTAAAAAGGACAAAAAGAAG
AAGGCTGATGAAACTCAAGCCTTACCGCAGAGACAGAAGAAACAGCAAACTGTGACTCTTCTT
CCTGCTGCAGATTTGGATGATTTCTCCAAACAATTGCAACAATCCATGAGCAGTGCTGACTCA
ACTCAGGCCTAAtaaaccggtaagacactgggtgtgaaccagcgcggcgagctgtgcgtccgtggccccatgatcatga
gcggctacgttaacaaccccgaggctacaaacgctctcatcgacaaggacggctggctgcacagcggcgacatcgcctactg
ggacgaggacgagcacttcttcatcgtggaccggctgaagagcctgatcaaatacaagggctaccaggtagccccagccgaa
ctggagagcatcctgctgcaacaccccaacatcttcgacgccggggtcgccggcctgcccgacgacgatgccggcgagctgc
ccgccgcagtcgtcgtgctggaacacggtaaaaccatgaccgagaaggagatcgtggactatgtggccagccaggttacaac
cgccaagaagctgcgcggtggtgttgtgttcgtggacgaggtgcctaaaggactgaccggcaagttggacgcccgcaagatcc

gcgagattctcattaaggccaagaagggcggcaagatcgccgtgtaaaggatccctccccccccccctaacgttactggccgaa
gccgcttggaataaggccggtgtgcgtttgtctatatgttattttccaccatattgccgtcttttggcaatgtgagggcccggaaacct
ggccctgtcttcttgacgagcattcctaggggtctttcccctctcgccaaaggaatgcaaggtctgttgaatgtcgtgaaggaagca
gttcctctggaagcttcttgaagacaaacaacgtctgtagcgaccctttgcaggcagcggaaccccccacctggcgacaggtgc
ctctgcggccaaaagccacgtgtataagatacacctgcaaaggcggcacaaccccagtgccacgttgtgagttggatagttgtg
gaaagagtcaaatggctctcctcaagcgtattcaacaaggggctgaaggatgcccagaaggtaccccattgtatgggatctgatc
tggggcctcggtgcacatgctttacatgtgtttagtcgaggttaaaaaaacgtctaggccccccgaaccacggggacgtggttttc
ctttgaaaaacacgatgataatatggccacacatatggcccagtccaagcacggcctgaccaaggagatgaccatgaagtacc
gcatggagggctgcgtggacggccacaagttcgtgatcaccggcgagggcatcggctaccccttcaagggcaagcaggccat
caacctgtgcgtggtggagggcggcccccttgcccttcgccgaggacatcttgtccgccgccttcatgtacggcaaccgcgtgttc
accgagtacccccaggacatcgtcgactacttcaagaactcctgccccgccggctacacctgggaccgctccttcctgttcgag
gacggcgccgtgtgcatctgcaacgccgacatcaccgtgagcgtggaggagaactgcatgtaccacgagtccaagttctacgg
cgtgaacttccccgccgacggccccgtgatgaagaagatgaccgacaactgggagccctcctgcgagaagatcatccccgtg
cccaagcagggcatcttgaagggcgacgtgagcatgtacctgctgctgaaggacggtggccgcttgcgctgccagttcgacac
cgtgtacaaggccaagtccgtgccccgcaagatgcccgactggcacttcatccagcacaagctgacccgcgaggaccgcag
cgacgccaagaaccagaagtggcacctgaccgagcacgccatcgcctccggctccgccttgccctgaatcgatagatcctaat
caacctctggattacaaaatttgtgaaagattgactggtattcttaactatgttgctccttttacgctatgtggatacgctgctttaatgcc
tttgtatcatgctattgcttcccgtatggctttcattttctcctccttgtataaatcctggttgctgtctctttatgaggagttgtggcccgttg
tcaggcaacgtggcgtggtgtgcactgtgtttgctgacgcaacccccactggttggggcattgccaccacctgtcagctcctttcc
gggactttcgctttccccctccctattgccacggcggaactcatcgccgcctgccttgcccgctgctggacaggggctcggctgtt
gggcactgacaattccgtggtgttgtcggggaaatcatcgtcctttccttggctgctcgcctgtgttgccacctggattctgcgcggg
acgtccttctgctacgtcccttcggccctcaatccagcggaccttccttcccgcggcctgctgccggctctgcggcctcttccgcg
tcttcgccttcgccctcagacgagtcggatctccctttgggccgcctccccgcctgagatcctttaagaccaatgacttacaaggc
agctgtagatcttagccactttttaaaagaaaaggggggactggaagggctaattcactcccaacgaagacaagatctgctttttg
cttgtactgggtctctctggttagaccagatctgagcctgggagctctctggctaactagggaacccactgcttaagcctcaataaa
gcttgccttgagtgcttcaagtagtgtgtgcccgtctgttgtgtgactctggtaactagagatccctcagacccttttagtcagtgtgg
aaaatctctagcagtagtagttcatgtcatcttattattcagtatttataacttgcaaagaaatgaatatcagagagtgagaggcccgg
gttaattaaggaaagggctagatcattcttgaagacgaaagggcctcgtgatacgcctatttttataggttaatgtcatgataataatg
gtttcttagacgtcaggtggcacttttcggggaaatgtgcgcggaacccctatttgtttatttttctaaatacattcaaatatgtatccgc
tcatgagacaataaccctgataaatgcttcaataatattgaaaaaggaagagtatgagtattcaacatttccgtgtcgcccttattccc
ttttttgcggcattttgccttcctgtttttgctcacccagaaacgctggtgaaagtaaaagatgctgaagatcagttgggtgcacgagt
gggttacatcgaactggatctcaacagcggtaagatccttgagagttttcgccccgaagaacgttttccaatgatgagcacttttaa
agttctgctatgtggcgcggtattatcccgtgttgacgccgggcaagagcaactcggtcgccgcatacactattctcagaatgactt
ggttgagtactcaccagtcacagaaaagcatcttacggatggcatgacagtaagagaattatgcagtgctgccataaccatgagt
gataacactgcggccaacttacttctgacaacgatcggaggaccgaaggagctaaccgcttttttgcacaacatgggggatcatg
taactcgccttgatcgttgggaaccggagctgaatgaagccataccaaacgacgagcgtgacaccacgatgcctgtagcaatg
gcaacaacgttgcgcaaactattaactggcgaactacttactctagcttcccggcaacaattaatagactggatggaggcggata
aagttgcaggaccacttctgcgctcggcccttccggctggctggtttattgctgataaatctggagccggtgagcgtgggtctcgc
ggtatcattgcagcactggggccagatggtaagccctcccgtatcgtagttatctacacgacggggagtcaggcaactatggatg
aacgaaatagacagatcgctgagataggtgcctcactgattaagcattggtaactgtcagaccaagtttactcatatatactttagat
tgatttaaaacttcatttttaatttaaaaggatctaggtgaagatcctttttgataatctcatgaccaaaatcccttaacgtgagttttcgtt
ccactgagcgtcagaccccgtagaaaagatcaaaggatcttcttgagatcctttttttctgcgcgtaatctgctgcttgcaaacaaa
aaaaccaccgctaccagcggtggtttgtttgccggatcaagagctaccaactctttttccgaaggtaactggcttcagcagagcgc
agataccaaatactgttcttctagtgtagccgtagttaggccaccacttcaagaactctgtagcaccgcctacatacctcgctctgct
aatcctgttaccagtggctgctgccagtggcgataagtcgtgtcttaccgggttggactcaagacgatagttaccggataaggcgc
agcggtcgggctgaacggggggttcgtgcacacagcccagcttggagcgaacgacctacaccgaactgagatacctacagc
gtgagctatgagaaagcgccacgcttcccgaagggagaaaggcggacaggtatccggtaagcggcagggtcggaacagga
gagcgcacgagggagcttccagggggaaacgcctggtatctttatagtcctgtcgggtttcgccacctctgacttgagcgtcgatt
tttgtgatgctcgtcaggggggcggagcctatggaaaaacgccagcaacgcggcctttttacggttcctggccttttgctggcctttt

gctcacatgttctttcctgcgttatcccctgattctgtggataaccgtattaccgcctttgagtgagctgataccgctcgccgcagccg
aacgaccgagcgcagcgagtcagtgagcgaggaagcggaagagcgcccaatacgcaaaccgcctctccccgcgcgttggc
cgattcattaatgcagcaagctcatggctgactaattttttttatttatgcagaggccgaggccgcctcggcctctgagctattccaga
agtagtgaggaggcttttttggaggcctaggcttttgcaaaaagctccccgtggcacgacaggtttcccgactggaaagcgggca
gtgagcgcaacgcaattaatgtgagttagctcactcattaggcaccccaggctttacactttatgcttccggctcgtatgttgtgtgg
aattgtgagcggataacaatttcacacaggaaacagctatgacatgattacgaatttcacaaataaagcatttttttcactgcattcta
gttgtggtttgtccaaactcatcaatgtatcttatcatgtctggatcaactggataactcaagctaaccaaaatcatcccaaacttccc
accccataccctattaccactgccaattacctgtggtttcatttactctaaacctgtgattcctctgaattattttcattttaaagaaattgt
atttgttaaatatgtactacaaacttagtagt

Vector name: VSV-G

ggatcccctgagggggcccccatgggctagaggatccggcctcggcctctgcataaataaaaaaaattagtcagccatgagctt
ggcccattgcatacgttgtatccatatcataatatgtacatttatattggctcatgtccaacattaccgccatgttgacattgattattga
ctagttattaatagtaatcaattacggggtcattagttcatagcccatatatggagttccgcgttacataacttacggtaaatggcccg
cctggctgaccgcccaacgacccccgcccattgacgtcaataatgacgtatgttcccatagtaacgccaatagggactttccatt
gacgtcaatgggtggagtatttacggtaaactgcccacttggcagtacatcaagtgtatcatatgccaagtacgccccctattgac
gtcaatgacggtaaatggcccgcctggcattatgcccagtacatgaccttatgggactttcctacttggcagtacatctacgtattag
tcatcgctattaccatggtgatgcggttttggcagtacatcaatgggcgtggatagcggtttgactcacggggatttccaagtctcca
ccccattgacgtcaatgggagtttgttttggcaccaaaatcaacgggactttccaaaatgtcgtaacaactccgccccattgacgc
aaatgggcggtaggcgtgtacggtgggaggtctatataagcagagctcgtttagtgaaccgtcagatcgcctggagacgccatc
cacgctgttttgacctccatagaagacaccgggaccgatccagcctcccctcgaagcttacatgtggtaccgagctcggatcctg
agaacttcagggtgagtctatgggacccttgatgttttctttccccttcttttctatggttaagttcatgtcataggaaggggagaagta
acagggtacacatattgaccaaatcagggtaattttgcatttgtaattttaaaaaatgctttcttcttttaatatactttttttgtttatcttatttc
taatactttccctaatctctttctttcagggcaataatgatacaatgtatcatgcctctttgcaccattctaaagaataacagtgataattt
ctgggttaaggcaatagcaatatttctgcatataaatatttctgcatataaattgtaactgatgtaagaggtttcatattgctaatagcag
ctacaatccagctaccattctgcttttattttatggttgggataaggctggattattctgagtccaagctaggcccttttgctaatcatgtt
catacctcttatcttcctcccacagctcctgggcaacgtgctggtctgtgtgctggcccatcactttggcaaagcacgtgagatctg
aattctgacactatgaagtgccttttgtacttagcctttttattcattggggtgaattgcaagttcaccatagtttttccacacaaccaaa
aaggaaactggaaaaatgttccttctaattaccattattgcccgtcaagctcagatttaaattggcataatgacttaataggcacagc
cttacaagtcaaaatgcccaagagtcacaaggctattcaagcagacggttggatgtgtcatgcttccaaatgggtcactacttgtg
atttccgctggtatggaccgaagtatataacacattccatccgatccttcactccatctgtagaacaatgcaaggaaagcattgaac
aaacgaaacaaggaacttggctgaatccaggcttccctcctcaaagttgtggatatgcaactgtgacggatgccgaagcagtgat
tgtccaggtgactcctcaccatgtgctggttgatgaatacacaggagaatgggttgattcacagttcatcaacggaaaatgcagca
attacatatgccccactgtccataactctacaacctggcattctgactataaggtcaaagggctatgtgattctaacctcatttccatg
gacatcaccttcttctcagaggacggagagctatcatccctgggaaaggagggcacagggttcagaagtaactactttgcttatg
aaactggaggcaaggcctgcaaaatgcaatactgcaagcattggggagtcagactcccatcaggtgtctggttcgagatggctg
ataaggatctctttgctgcagccagattccctgaatgcccagaagggtcaagtatctctgctccatctcagacctcagtggatgtaa
gtctaattcaggacgttgagaggatcttggattattccctctgccaagaaacctggagcaaaatcagagcgggtcttccaatctctc
cagtggatctcagctatcttgctcctaaaaacccaggaaccggtcctgctttcaccataatcaatggtaccctaaaatactttgaga
ccagatacatcagagtcgatattgctgctccaatcctctcaagaatggtcggaatgatcagtggaactaccacagaaagggaact
gtgggatgactgggcaccatatgaagacgtggaaattggacccaatggagttctgaggaccagttcaggatataagtttcctttata
catgattggacatggtatgttggactccgatcttcatcttagctcaaaggctcaggtgttcgaacatcctcacattcaagacgctgctt
cgcaacttcctgatgatgagagtttattttttggtgatactgggctatccaaaaatccaatcgagcttgtagaaggttggttcagtagtt
ggaaaagctctattgcctctttttctttatcatagggttaatcattggactattcttggttctccgagttggtatccatctttgcattaaatta
aagcacaccaagaaaagacagatttatacagacatagagatgaaccgacttggaaagtaactcaaatcctgcacaacagattct
tcatgtttggaccaaatcaacttgtgataccatgctcaaagaggcctcaattatatttgagtttttaatttttatgaaaaaaaaaaaaaa
aaacggaattcaccccaccagtgcaggctgcctatcagaaagtggtggctggtgtggctaatgccctggcccacaagtatcact
aagctcgctttcttgctgtccaatttctattaaaggttcctttgttccctaagtccaactactaaactgggggatattatgaagggccttg

agcatctggattctgcctaataaaaaacatttattttcattgcaatgatgtatttaaattatttctgaatattttactaaaaagggaatgtgg
gaggtcagtgcatttaaaacataaagaaatgaagagctagttcaaaccttgggaaaatacactatatcttaaactccatgaaagaa
ggtgaggctgcaaacagctaatgcacattggcaacagcccctgatgcctatgccttattcatccctcagaaaaggattcaagtag
aggcttgatttggaggttaaagttttgctatgctgtattttacattacttattgttttagctgtcctcatgaatgtcttttcactacccatttgct
tatcctgcatctctcagccttgactccactcagttctcttgcttagagataccacctttcccctgaagtgttccttccatgttttacggcg
agatggtttctcctcgcctggccactcagccttagttgtctctgttgtcttatagaggtctacttgaagaaggaaaaacaggggggcat
ggtttgactgtcctgtgagcccttcttccctgcctcccccactcacagtgacccggaatccctcgacatggcagtctagcactagtg
cggccgcagatctgcttcctcgctcactgactcgctgcgctcggtcgttcggctgcggcgagcggtatcagctcactcaaaggc
ggtaatacggttatccacagaatcaggggataacgcaggaaagaacatgtgagcaaaaggccagcaaaaggccaggaaccg
taaaaaggccgcgttgctggcgtttttccataggctccgcccccctgacgagcatcacaaaaatcgacgctcaagtcagaggtg
gcgaaacccgacaggactataaagataccaggcgtttccccctggaagctccctcgtgcgctctcctgttccgaccctgccgctt
accggatacctgtccgcctttctcccttcgggaagcgtggcgctttctcatagctcacgctgtaggtatctcagttcggtgtaggtcg
ttcgctccaagctgggctgtgtgcacgaaccccccgttcagcccgaccgctgcgccttatccggtaactatcgtcttgagtccaac
ccggtaagacacgacttatcgccactggcagcagccactggtaacaggattagcagagcgaggtatgtaggcggtgctacaga
gttcttgaagtggtggcctaactacggctacactagaagaacagtatttggtatctgcgctctgctgaagccagttaccttcggaaa
aagagttggtagctcttgatccggcaaacaaaccaccgctggtagcggtggtttttttgtttgcaagcagcagattacgcgcagaa
aaaaaggatctcaagaagatcctttgatcttttctacggggtctgacgctcagtggaacgaaaactcacgttaagggattttggtcat
gagattatcaaaaaggatcttcacctagatccttttaaattaaaaatgaagttttaaatcaatctaaagtatatatgagtaaacttggtct
gacagttaccaatgcttaatcagtgaggcacctatctcagcgatctgtctatttcgttcatccatagttgcctgactccccgtcgtgta
gataactacgatacgggagggcttaccatctggccccagtgctgcaatgataccgcgagacccacgctcaccggctccagattt
atcagcaataaaccagccagccggaagggccgagcgcagaagtggtcctgcaactttatccgcctccatccagtctattaattgt
tgccgggaagctagagtaagtagttcgccagttaatagtttgcgcaacgttgttgccattgctacaggcatcgtggtgtcacgctcg
tcgtttggtatggcttcattcagctccggttcccaacgatcaaggcgagttacatgatcccccatgttgtgcaaaaaagcggttagct
ccttcggtcctccgatcgttgtcagaagtaagttggccgcagtgttatcactcatggttatggcagcactgcataattctcttactgtc
atgccatccgtaagatgcttttctgtgactggtgagtactcaaccaagtcattctgagaatagtgtatgcggcgaccgagttgctctt
gcccggcgtcaatacgggataataccgcgccacatagcagaactttaaaagtgctcatcattggaaaacgttcttcggggcgaa
aactctcaaggatcttaccgctgttgagatccagttcgatgtaacccactcgtgcacccaactgatcttcagcatcttttactttcacc
agcgtttctgggtgagcaaaaacaggaaggcaaaatgccgcaaaaaagggaataagggcgacacggaaatgttgaatactca
tactcttcctttttcaatattattgaagcatttatcagggttattgtctcatgagcggatacatatttgaatgtatttagaaaaataaacaaa
taggggttccgcgcacatttccccgaaaagtgccacctgacgt
